# Affinity and distance dependence of SLiM-mediated dephosphorylation by protein phosphatase 1

**DOI:** 10.64898/2026.09.09.750541

**Authors:** Evi Setiani Lande, Magnus Kjaergaard

## Abstract

Protein phosphatases counterbalance kinases by dephosphorylating phospho-proteins to regulate signaling pathways. But unlike kinases, their shallow catalytic groove has limited selectivity for the phospho-peptide motifs. Phosphatases such as protein phosphatase-1 (PP1) recognize their substrates through short linear motifs (SLiMs) within intrinsically disordered regions, yet the mechanism of SLiM-mediated recruitment remains unclear. We developed an intrinsically disordered phospho-protein substrate to enable quantitative modelling of tethered PP1 catalysis by varying SLiM affinity and distance. We show that a PP1-binding SLiM linked to a phospho-peptide by a flexible spacer is sufficient to enhance dephosphorylation. Dephosphorylation occurs most efficiently at a spacing of 20 – 30 residues, in agreement with predictions from theoretical polymer models. Low micromolar dissociation constants are most efficient for dephosphorylation, which can be modelled numerically as a trade-off between substrate binding and substrate inhibition. Bivalent SLiMs enhance or decrease catalysis depending on the combined affinity. Together, these results define a quantitative and generalizable framework for how linear motifs direct dephosphorylation by PP1.

## Introduction

Protein phosphorylation regulates protein interactions, stability and activity. Phosphorylation levels reflect a tug-of-war between enzymes that add phosphate groups – kinases, and enzymes that remove phosphate groups – phosphoprotein phosphatases (PPP). Phosphoprotein phosphatases such as protein phosphatase-1 (PP1) exhibit limited intrinsic substrate selectivity due to the shallow nature of its catalytic cleft [1]. Initially, this led to the assumption that kinases were the primary determinants of phosphorylation levels, with phosphatases such as PP1 passively resetting phosphorylation levels. However, substrate profiling has revealed that phosphatases have distinct substrate preferences in cells [2] and *in vitro* [3], and that their dysfunction is a key pathological mechanism [4,5]. Phosphatases and kinases are thus equal players in regulating phosphorylation levels although they achieve their specificity in different ways.

Phosphatase profiling has revealed that their substrate selectivity primarily depends on short linear motifs (SLiMs) distal from the phospho-site. These motifs typically consist of four to ten residues located within intrinsically disordered regions (IDRs) of regulatory partners or direct substrates. In PP1, such motifs include RVxF, SILK, ΦΦ, KiR, and MyPhoNE [6]. The SLiMs are recognized by binding grooves on the phosphatase and they guide the substrate to the active site. Proteomic studies have shown that the SLiM may be thousands of residues away from the phospho-site [2,3]. Despite the central role of SLiMs in PP1 substrate recognition, there is currently no quantitative model for how SLiMs regulate dephosphorylation, and it is thus unclear how activity depends on, for example the affinity of the SLiM-phosphatase interaction, the SLiM’s position relative to the phospho-site, the nature of the protein or protein complex, that bridge the SLiM and the phospho-site.

SLiM-mediated catalysis has been studied in other enzyme systems, providing a useful conceptual framework for phosphatases. In the simplest mechanism, the SLiM anchors the substrate to the enzyme and increases the effective concentration of substrate experienced by the active site. In this model, the segment between the SLiM and the phospho-site provides a flexible linker that allows enzyme and substrate to meet, but there are few constraints on the linker sequence beyond flexibility. The linker needs to be sufficiently long to span the distance between the SLiM docking pocket and the active site. Beyond this length, the effective concentrations is expected to decrease with linker length following a polymer scaling law [7,8]. In an engineered system, protein kinase A (PKA) was tethered to its substrate with linkers of variable length, which showed that the intra-complex phosphorylation depends on the effective concentration enforced by the linker [9]. For steady-state kinetics, the catalytic rates showed a bell-shaped dependence on the affinity of the docking interaction, which can be explained by a trade-off between limiting substrate binding and product inhibition [10]. We hypothesize that similar principles govern SLiM-mediated dephosphorylation kinetics by PP1.

Here, we develop a modular PP1 substrate system where SLiM affinity and spacing can be independently controlled, thus providing a degree of control unachievable in natural substrates. We show that tethering a phospho-peptide to the enzyme via a disordered linker is sufficient to enhance its dephosphorylation rate and that the optimal spacing between SLiM and phospho-site is 20 – 30 residues. We vary the affinity of the docking interaction through mutagenesis and multivalency, which suggests an optimal affinity in the low micromolar range. Comparison to numerical simulations using a theoretical model for tethered catalysis broadly agrees with the optimal affinity for optimal dephosphorylation, however the affinity range where the reaction rate is enhanced was measured to be narrower than suggested by the model. In summary, the results are consistent with a mechanism where SLiMs enhance dephosphorylation by increasing the effective concentration of the substrate. Together, these findings provide a theoretical model for SLiM-mediated dephosphorylation.

## Results

### SLiMs facilitate tethered catalysis

To understand the role of SLiMs in PP1 catalysis, we developed synthetic substrates consisting of a SLiM (RVAF) connected to the PKA substrate, kemptide [11], by a flexible disordered linker of variable length (Fig. 1A). The two arginine residues at positions -2 and -3 relative to the phosphoserine in kemptide (LRRASLG) match the preferred substrate motif of PP1 [3]. To aid purification, thioredoxin-1 (Trx-1) and a His_6_-tag were fused to the N-terminus and a StrepII-tag [12] at the C-terminus to allow sequential affinity purification. The purified substrates (Fig. S1) were quantitatively phosphorylated using PKA as confirmed by Phos-tag gels [13] (Fig. 1A). When the phospho-acceptor serine was substituted with alanine, no mobility shift was seen upon PKA incubation (Fig. 1A, S2) showing that phosphorylation only happens at the intended site.

**Fig. 1.**
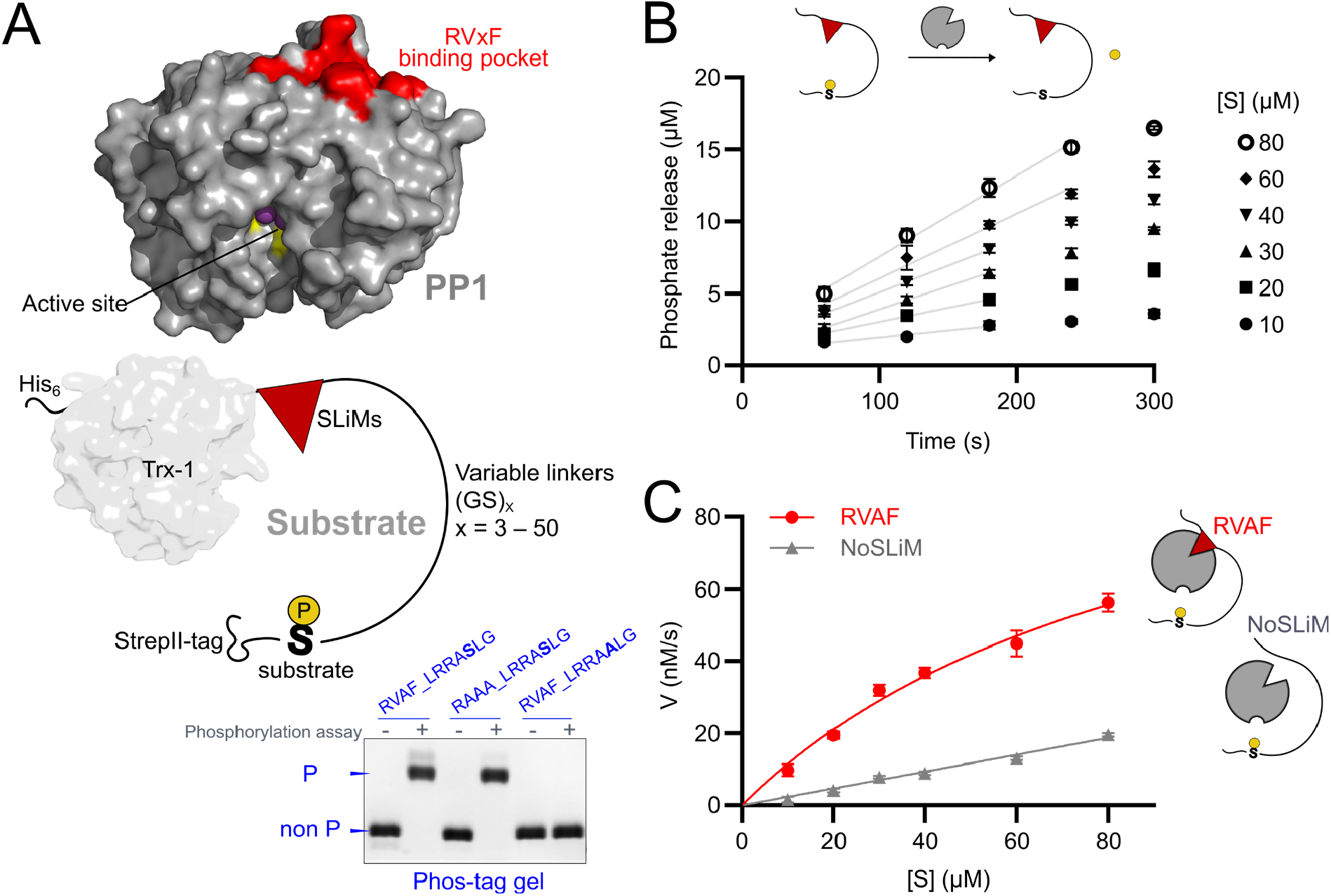
A model system for studying SLiM-tethered dephosphorylation kinetics. **A)** Schematic representation of PP1 using PyMOL, highlighting the docking pocket for SLiM (red residues) and an active site for dephosphorylation (yellow residues with 2 purple spherical of Mn^2+^ ions). Synthetic disordered substrate construct consists of a SLiM, disordered linker (GS repeats), and a phospho-site (kemptide), with His_6_-tag and Thioredoxin-1 (Trx-1) as a soluble domain at the N-terminus and StrepII-tag at C-terminus. Linear disordered linkers are varied from 6 to 100 residues. The Phos-tag gel displays substrate phosphorylation status of representative full-length substrates. **B)** Phosphate release of RVAF_(GS)_10__kemptide measured using malachite green assay across different substrate concentrations. Grey lines are linear regressions. **C)** Comparison of dephosphorylation kinetics between RVAF substrate variant (red) and NoSLiM or untethered (grey) substrate variants with (GS)_10_ spacer. Error bars indicate standard error of means of the linear regression.

To assess the effect of SLiMs, we compared the dephosphorylation of phospho-peptide substrates with either a consensus RVxF SLiM joined by a 20-residue linker or a substrate without a SLiM. The enzymatic activity was measured using malachite green assay that allows the rate of phosphate release to be determined for micromolar substrate concentrations (Fig. 1B, S3). In the absence of a binding motif (NoSLiM), dephosphorylation rate increases linearly with concentration without signs of saturation (Fig. 1C). In addition to that, the NoSLiM substrate has approximately twice as high maximum rate (V_max_) than SLiM-tethered variants. The substrate with the SLiM has increased dephosphorylation rates and a concentration dependence indicative of saturation in the mid-micromolar range (Fig. 1C). This showed that no particular sequence features in the substrate dephosphorylated by PP1 beyond the phospho-site. No phosphate release was detected for a substrate without the phospho-serine (Fig. S3C). This substrate provides a robust model for quantifying how SLiMs regulate PP1-dependent dephosphorylation.

### Disordered linkers modulate the kinetics through distance-dependent tethering effects

Phosphatase substrates show a variety of spacings between the enzyme targeting SLiMs and the phospho-site [6], which suggests few structural constraints for forming a catalytically competent complex. We modelled the simplest scenario of a phospho-site connected to RVxF motif through a disordered linker. The worm-like chain model [8] predicts end-to-end probability distributions (p(r)) for each linker length and a characteristic upper reach of the linker (Fig. 2A). The linkers were modelled with a persistence length of 4 Å, which reproduce the compaction of other disordered linkers [14]. We estimated the separation between the RVxF-binding pocket and the PP1 active site to be ∼27 Å based on the crystal structure (PDB: 3E7A) [15] (Fig. 2B). Using this distance as the distance from anchor points, the worm-like chain model predicts C_eff_ as a function of linker length (Fig. 2C). The model predicts that spacings of less than ∼10 residues are insufficient with bivalent binding, whereas a 23-residue spacing provides the optimal effective concentration. For longer linkers, the effective concentration gradually declines. Together, these calculations provide a quantitative prediction for how the spacing between the SLiM and the phospho-site affects dephosphorylation efficiency.

**Fig. 2.**
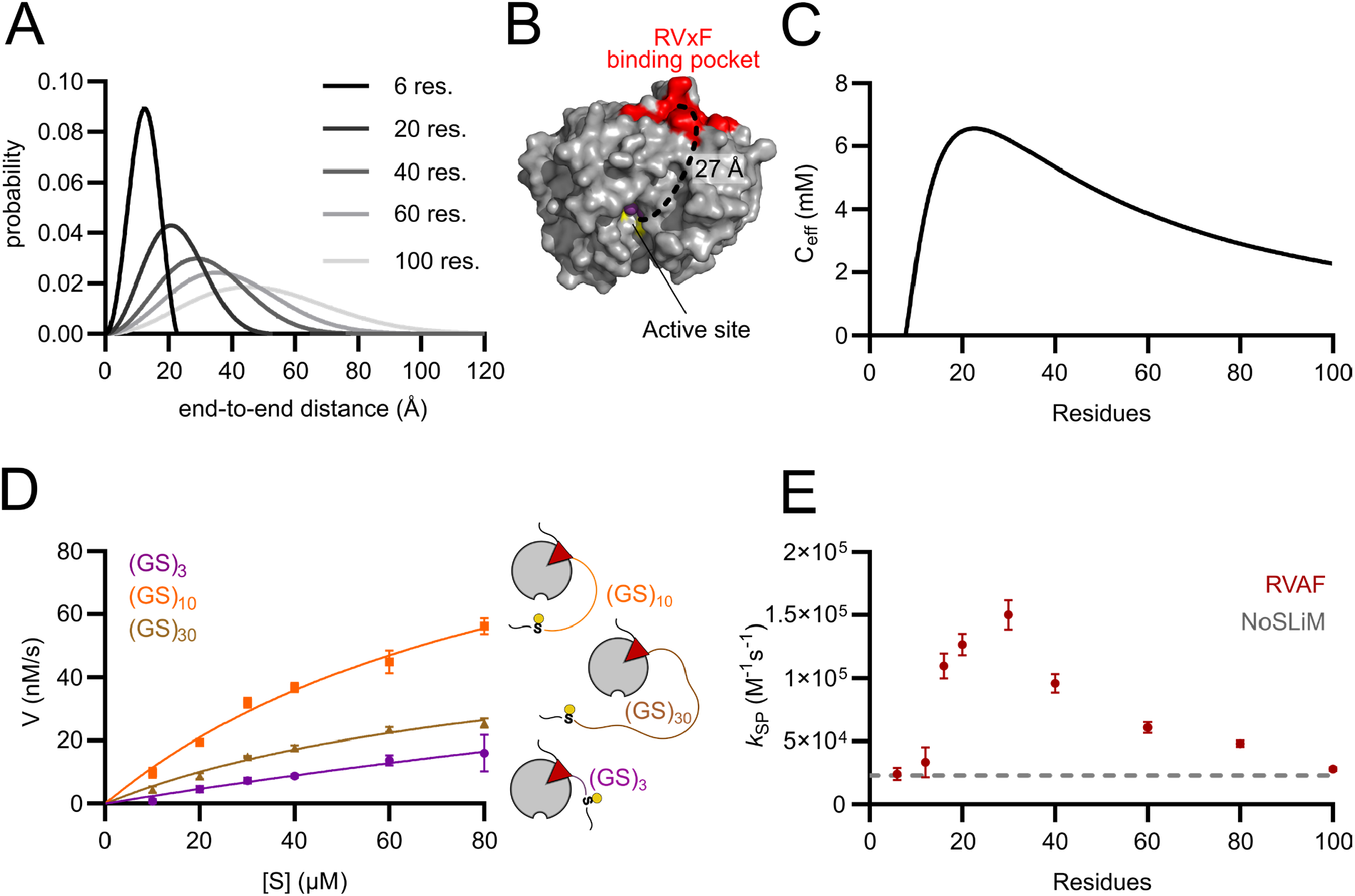
Linker-length dependence kinetics of dephosphorylation by PP1. **A)** Simulated end-to-end distance probability distribution p(r) of 6 to 100 residues linear linker. **B)** Distance measured between RVxF docking pocket (red) and the active site (yellow) using PyMOL. **C)** Simulated effective concentration C_eff_ plotted as a function of linker length with distance of 27Å from RVxF docking pocket to the active site. **D)** Comparison of dephosphorylation kinetics of RVAF substrates with 6, 20 and 60-residue linkers. **E)** Specificity constant *k*_SP_ comparison of RVAF_kemptide with various linker lengths from 6 to 100 residues (red). Untethered substrate variant (NoSLiM) is displayed in grey dash as baseline. Figure 2A and 2C are made using C_eff_ calculator (https://chemeslab.myqnapcloud.com:3939/CeffApp/). Error bars show standard error of means of the specificity constant (*k*_SP_) equation fitting.

To test this prediction, we measured *in vitro* dephosphorylation kinetics across a range of linker lengths. We made substrates with disordered linkers ranging from 6 to 100 residues that are both above and below the predicted optimal spacing. For each variant, we measured dephosphorylation rate as a function of substrate concentration (Fig. 2D, S4A). For the shortest linker, dephosphorylation rates were almost linear with substrate concentration with a similar rate to a NoSLiM substrate, confirming that there is a minimum threshold for spacing between the SLiM and the phospho-site. In contrast, longer linkers showed elevated dephosphorylation rates and concentration-dependent saturation.

To quantitatively compare substrates at their initial concentration without reaching saturation, we fitted the catalytic rates to the Michaelis-Menten equation expressed via the specificity constant (*k*_SP_) [16] corresponding to the catalytic efficiency (*k*_cat_/*K*_M_). Overall, the linker length dependence of *k*_SP_ (Fig. 2E) resembles the C_eff_ curve predicted by the worm-like chain model (Fig. 2C). The NoSLiM substrate defines the intrinsic catalytic efficiency as a baseline. Linkers up to 12 residues do not enhance dephosphorylation rates, whereas a 16-residue linker does. This effectively defines the minimum required spacing between SLiM and phospho-site. Optimal dephosphorylation rates are achieved by substrates with 20 – 30 residues linkers, in agreement with the prediction by the worm-like chain model. Beyond the optimal spacing, the catalytic efficiency dropped with increasing linker length. The decrease is steeper than predicted from the worm-like chain model and reaches the NoSLiM baseline at 100 residues – seemingly defining an upper limit for spacing between SLiM and phospho-site. The limited solubility of long GS-repeats prevented the series from being extended further. Overall, these results confirm that linker length critically determines tethered catalytic efficiency and defines optimal and lower bounds for the spacing between SLiM and phospho-site.

### A higher catalytic enhancement is provided by composite SLiMs

PP1 has several distinct docking pockets that bind motifs including SILK, MyPhoNE, and ΦΦ (Fig. 3A). These SLiMs are widely distributed in PP1 interactomes and often appear as composite interactions [14, 15]. Using the worm-like chain, we predicted C_eff_ for phospho-site to SLiM spacings for each motif (Fig. 3B), which indicated optimal spacings of 44, 19 and 9 residues for SILK, ΦΦ [19,20], and MyPhoNE [18] respectively. We tested each of these motifs in a similar model substrate as used for RVxF motif, using 20- and 40-residue linkers to accommodate the different spacings of docking sites (Fig. 3C). *In vitro* phosphate release rates of modified SLiMs are provided in Fig. S4B. The *k*_SP_ of SILK motif showed a slight enhancement compared to the NoSLiM control, whereas MyPhoNE and ΦΦ motifs were indistinguishable from baseline (Fig. 3D). This indicated that these individual motifs were not sufficient to target the substrate for PP1 catalysis.

**Fig. 3.**
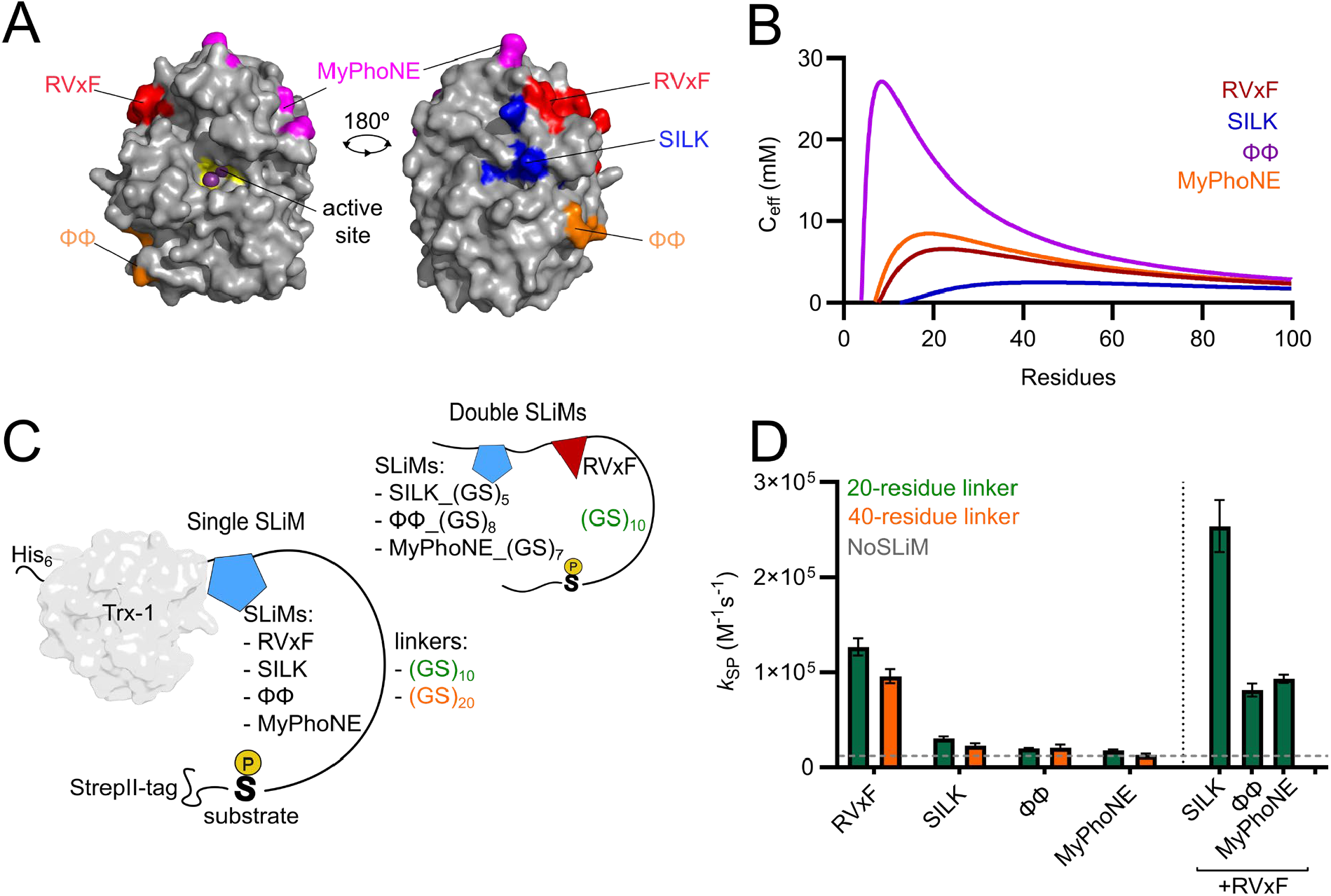
Developing a model for composite SLiMs. **A)** Different SLiM docking pockets in PP1 and their relative position to the active site. The binding sites were identified using crystal structure with PDBid: 7T0Y and 6CZO for RVxF, SILK and ΦΦ [19,20] and 1S70 for MyPhoNE [18]. **B)** Effective concentration C_eff_ plotted as a function of linker length with distance of 27Å, 37 Å, 25 Å, and 17 Å, respectively for RVxF, SILK, MyPhoNE and ΦΦ from their docking pocket to the active site. **C)** Schematic of substrates with individual SLiMs and their RVxF composite motifs. The linker length (GS)_x_ was adjusted to fit the distance from each docking pocket to RVxF’s (right panel). **D)** Comparison of substrate’s specificity constant *k*_SP_ of single SLiM with 20 and 40-residue linkers on the left side and double SLiMs with 20-residue linker from RVxF to phospho-site on the right side. Error bars show standard error of means of the *k*_SP_.

SLiMs often form multivalent interactions involving several SLiMs. Next, we tested whether these motifs could synergize with another motif to enhance dephosphorylation by combining RVxF with secondary motifs connected by a linker that match the distances between their binding pockets. Among all combinations tested, only the SILK+RVxF composite increased *k*_SP_ beyond that of RVxF alone (Fig. 3D). In contrast, other combinations, which are MyPhoNE+RVxF and ΦΦ+RVxF, yielded slightly lower *k*_SP_ (Fig. 3D). However, reduced catalytic activity does not necessarily reflect weaker binding, which thus led us to compare both binding affinities.

### SLiM affinity defines an optimal window for catalytic efficiency

To assess affinity–activity relationship, we measured dissociation constants of docking motifs using a fluorescence polarization (FP) competition assay. We used a fluorescein isothiocyanate (FITC)-labelled 15-residue peptide from the nuclear PP1 interactor NIPP1 [21] as the probe, which has an extended binding site centered at the RVxF pocket (Fig. 4A). Direct titration with FP yielded a *K*_d_ of 252 nM (Fig 4B) in agreement with previous studies measured using ITC and a high-throughput *in vitro* screening [2,19]. To avoid off target phosphorylation on RVTF’s threonine, we used non-phosphorylatable T202A variants in substrates, whereas the higher affinity WT sequence was used in the competitor. FITC-labelled NIPP1 can be displaced by unlabeled RVxF-containing substrates (Fig. 4C) allowing the *K*_i_ or *K*_d_ to be extracted from the IC_50_ [22]. Surprisingly, the NoSLiM substrate also competed with NIPP1 at very high concentrations (Fig. 4C).

**Fig. 4.**
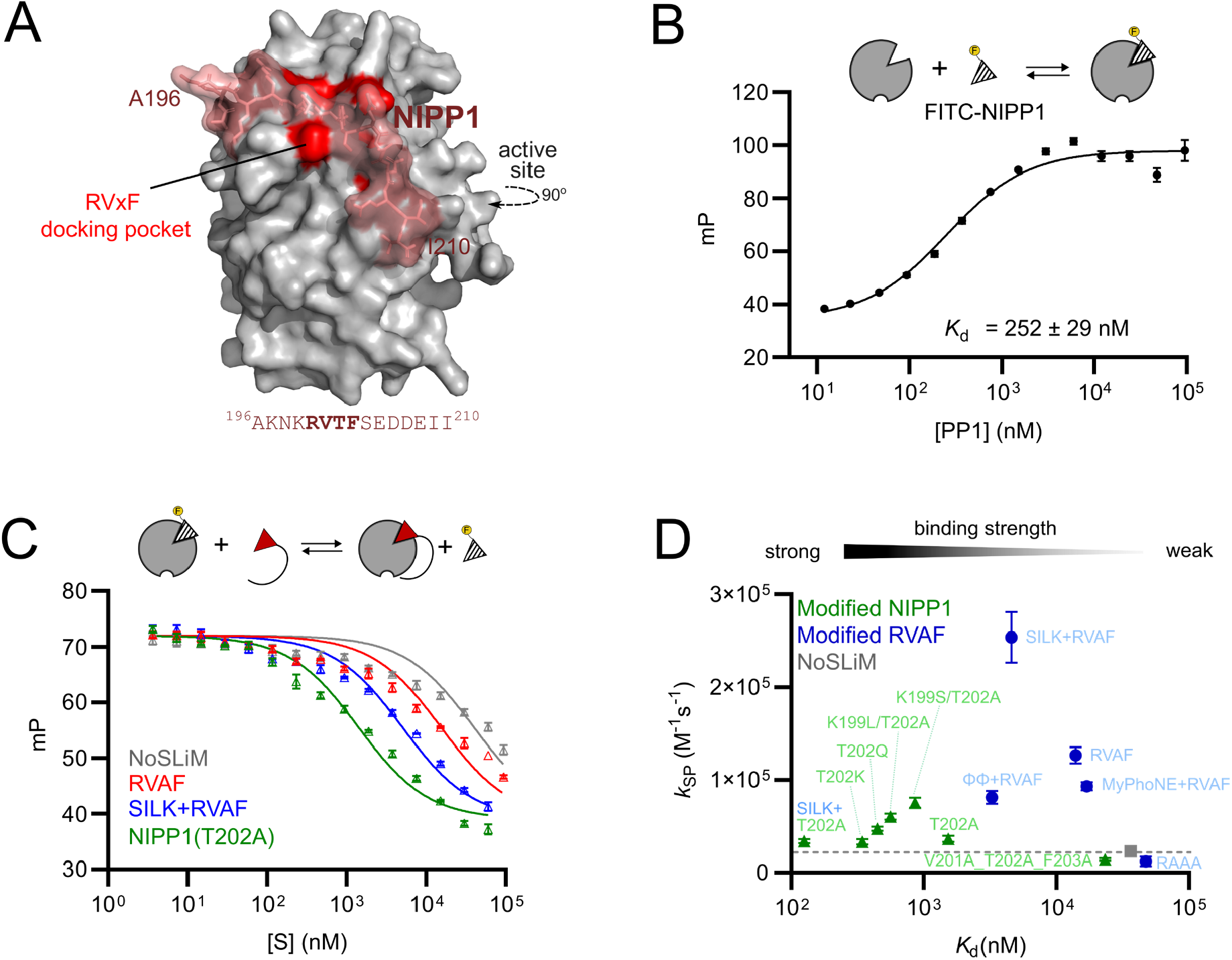
Affinity dependence of dephosphorylation kinetics by PP1. **A)** Nuclear inhibitor of protein phosphatase-1 (NIPP1)^196-210^ peptide docks on RVxF docking pocket (red) of PP1 (PDB: 3V4Y) [21]. **B)** Fluorescence polarization direct binding assay of FITC-labelled NIPP1 peptide and PP1 using 30 nM FITC-labelled NIPP1 peptide. **C)** Fluorescence-polarization competition assay of substrates across SLiMs. Titration was performed using 30 nM of FITC-labelled NIPP1 peptide and 250 nM of PP1. **D)** Specificity constants (*k*_SP_) plotted as a function of dissociation constant (*K*_d_) of modified RVxF substrates. Green data points are modified NIPP1-based substrates with one composite variant (SILK+T202A). Blue dots are RVAF-based substrate variants: RVAF, RAAA, and three composites: SILK+RVAF, ΦΦ+RFAV, and MyPhoNE+RVAF. Error bars show standard error of means of the *k*_SP_ equation fitting. Grey data point and dash line showing the *k*_SP_ of NoSLiM variant.

We evaluated substrate affinities with mono- and bivalent SLiMs using the FP competition assay (Fig. S5). The RVAF motif alone has a *K*_d_ of 14 µM, whereas it decreases to 3 – 5 µM in combination with the SILK or ΦΦ motifs. In contrast, adding MyPhoNE motif decreases the binding strength, which could correspond to its ability to form helical structure and thus introduce stickiness to the construct [18]. To extend the range of affinities, we created a new series of substrates containing the 15-residue motif from NIPP1 linked to kemptide. We tested a range of mutations previously found in NIPP1 to increase affinity in high through-put assays [3], including T202Q, K199L/T202A, K199S/T202A as well as a NIPP1 with a disrupted motif (V201A/R202A/F203A) and a combination of NIPP1 and SILK. NIPP1(T202A) peptide shared the same *K*_d_ with the substrate form with a kemptide embedded (Table S1), indicates that kemptide does not cause a binding affinity shift when the SLiM has much tighter affinity. Together, these two series span docking affinities across three orders of magnitude (Fig. S5).

We assessed the dephosphorylation kinetics of substrates with each of the NIPP1 variants and extracted the specificity constant for each variant (Fig. S6). NIPP1 SLiMs only increased dephosphorylation rates moderately with *k*_SP_ values up to ∼2-fold above NoSLiM. The relationship between SLiM binding affinity and catalytic efficiency can be evaluated by plotting *k*_SP_ against *K*_d_ of the docking motif (Fig. 4D). The relation between affinity and catalytic efficiency is non-monotonous bell-shaped curve, indicating that increasing affinity will increase catalytic efficiency up to a certain point beyond which increased affinity becomes inhibitory. The highest dephosphorylation rate for our substrates is observed at low micromolar SLiM affinity, approximately 4.5 µM, corresponding to the SILK+RVAF composite motif (Fig. 4D). Thus, optimal catalytic activity occurs within an intermediate affinity range. This broadly resembles the Sabatier principle in heterogeneous catalysis [23] and the pattern observed for tethering of kinases [10], which provides a theoretical framework for interpreting our results.

### A theoretical model for SLiM-mediated dephosphorylation

We sought to establish a kinetic model for the dependence of catalytic rates in SLiM affinity based on previous work in kinases [10]. Phospho-proteins without SLiMs are dephosphorylated at a basal rate, which is enhanced in the presence of a SLiM. Mechanistically, this suggests that dephosphorylation proceeds through two distinct pathways: untethered and SLiM-tethered (Fig. 5A). The untethered mechanism follows a Michaelis-Menten model, whereas SLiM-tethered catalysis involves initial recruitment to PP1 via SLiM binding. Ternary complexes can also occur (Fig. S8A), which are included in the simulations but omitted from Fig. 5A for clarity. In the tethered complex, the dephosphorylation rate depends on the linker between the SLiM and the phospho-site, which can be modelled using the effective concentration [9]. Crucially, the SLiM-tethered pathway requires dissociation of the docking interaction, which introduces the possibility of product inhibition. This represents a minimal reaction mechanism for tethered catalysis and can be simulated numerically from the rate constants of the individual steps.

**Fig. 5.**
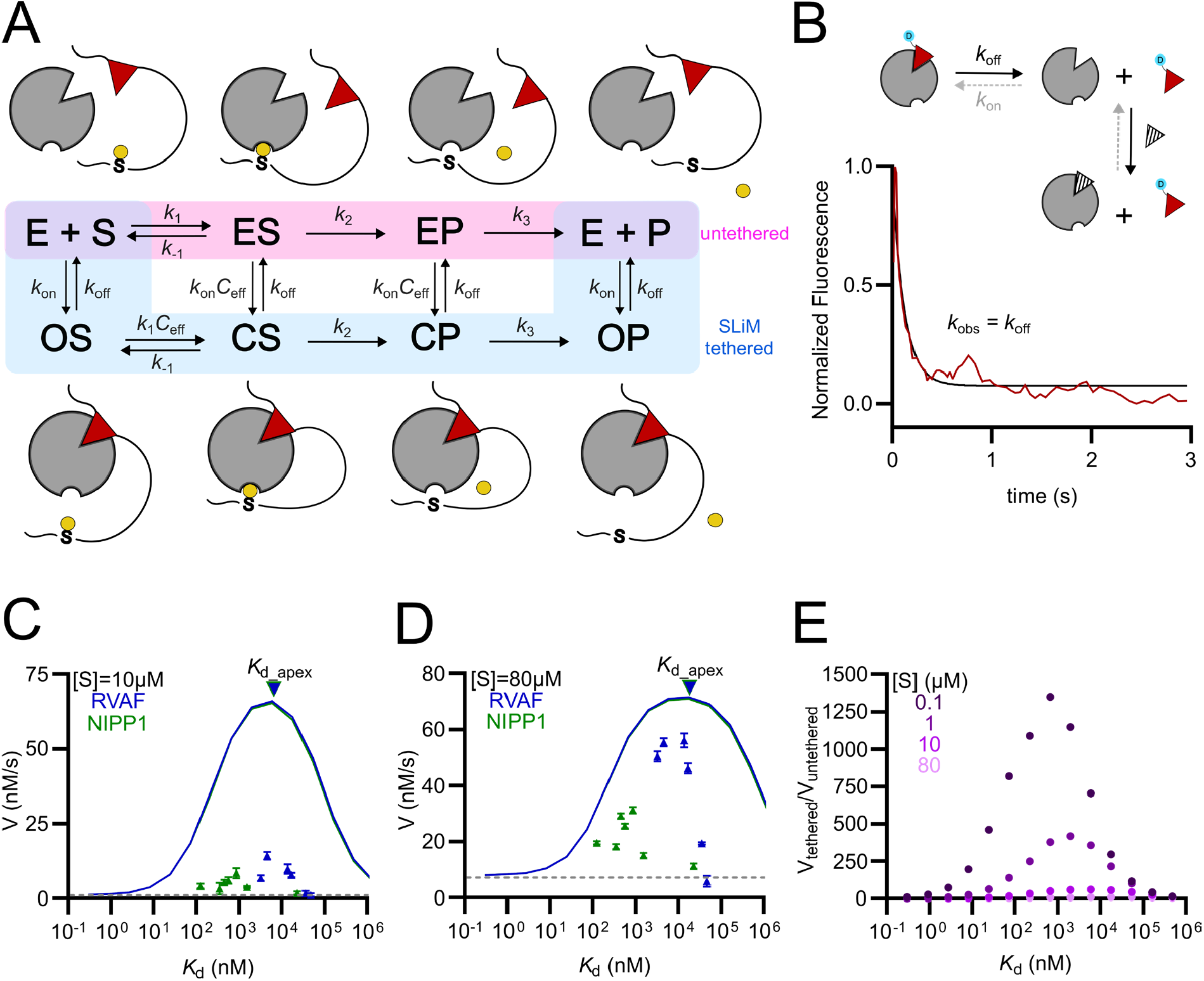
Numerical simulation of reaction rates in different substrate concentrations. **A)** Kinetic scheme of tethered and untethered reaction. Enzyme (E), substrate (S), and product (P) are shown. ES and EP indicate the enzyme’s active site is bound with the substrate or product. OS and OP denote open state of enzyme-substrate or enzyme-product complex, while CS and CP represent closed conformation of the complex. **B)** Schematic representation of mechanistic displacement of dansyl-labelled NIPP1 from PP1 by excess unlabelled peptide. The equilibrium constant favors the complex to dissociate due to excessive amount of competitor, resulting in free dansyl-labelled NIPP1 species. The observed kinetic rate thus corresponds to the dissociation rate of dansyl-labelled NIPP1 and PP1 (*k*_off_) of 8.4 ± 0.8 s^-1^. C – D) Numerical simulations (lines) and experimental data (triangles) of phosphate release rate as a function of *K*_d_ at substrate concentrations of 10 (C) and 80 µM (D). Blue features represent RVxF and green features represent NIPP1. Error bars show standard error of means of the dephosphorylation rate (V). Dash lines are untethered catalytic rates of each substrate concentration. Inverted triangles on the top of lines are optimum dissociation constant, *K*_d_apex_. **E)** Ratio of tethered and untethered phosphate release rate across extensive lower substrate concentrations from 0.1 – 80 µM derived from numerical simulations.

To obtain rate constants for the SLiM association, we performed stopped-flow measurements between PP1 and an environmentally sensitive dansyl-labelled NIPP1 peptide. The fluorescence trace of the association reaction had two phases with opposing fluorescence amplitudes (Fig. S7). The biphasic trace suggests the presence of at least two different bound conformations but precludes direct determination of *k*_on_. Instead, we measured *k*_off_ by kinetic displacement of dansyl-NIPP1 by a large excess of unlabeled NIPP1 (Fig. 5B) resulting in *k*_off_ of 8.4 s^-1^. In combination with the *K*_d_ from equilibrium studies, this allowed us to estimate a *k*_on_ of 3.3 × 10^7^ M^-1^ s^-1^, which is near the diffusion limit as seen for many interactions of disordered peptides [24]. As the difference in affinity of peptide interactions mainly arise from variations in *k*_off_, we used this *k*_on_ value for all substrate variants to extrapolate a *k*_off_ value from *K*_d_ for each variant for kinetic modeling. Many of the microscopic rate constants in the model presented in Fig. 5A are intractable to obtain experimentally and we thus used reasonable estimations based on available data and similar systems (details in Materials and Methods).

To examine how the catalytic rate varies with SLiM affinity, we simulated phosphate release numerically across the two primary pathways (Fig. 5A) and additional ternary complexes (Fig. S8A) while systematically varying *k*_off_. Simulations were performed at the lowest (Fig. 5C) and highest (Fig. 5D) substrate concentration tested experimentally. Both approaches suggest a *K*_d_-dependent bell-shaped phosphate release rate with an apex consistent with that predicted by an analytical equation [10]. At substrate concentration of 10 µM, the experimental phosphate release rates were ∼5 folds lower than those predicted by simulation (Fig. 5C). Nevertheless, the shape of the curve and the predicted apex were similar to that observed experimentally, at ∼6 µM *K*_d_ corresponding to the SILK+RVAF variant. At substrate concentration of 80 µM, the magnitude of simulated and experimental phosphate release rate was similar – in addition to the apex of the curve (Fig. 5D). The simulations differ systematically from experimental data in the width of the bell-shaped curve, where simulations suggest that dephosphorylation should be enhanced by a broader range of SLiM affinities – especially weaker SLiMs at 80 µM. This suggests that there is an additional contribution in the system not accounted for by the model. Overall, simulated phosphate release rates accurately captured the experimental trends. Importantly, both modeling and experiments support the existence of an optimal dissociation rate for efficient catalysis.

The physiological concentrations of most phospho-proteins are much lower than the substrate concentrations used in our experiments. The limits of quantification of the malachite green assay prevents us from conducting experiments at lower substrates concentrations, however the agreement between numerical simulation and experiments encouraged us to simulate experimentally intractable concentrations. To compare different substrates concentrations, we normalized the rates by the untethered rate at the same substrate concentration (Fig. 5E). This ratio quantifies the enhancement provided by a SLiM, which is relevant for the specificity relative to off-target substrates. At substrate concentrations of 1 and 0.1 µM, the model predicts rate enhancement of up to 400-fold and 1300-fold – much higher than 9-fold enhancement predicted at the highest concentration tested of experimentally observable high concentrations at 80 µM. Lower concentrations shifted the *K*_d_ to the maximal enhancement of sub-micromolar *K*_d_ values. In combination, these results suggest that SLiMs enhance dephosphorylation by increasing the effective concentration of the phospho-site at the PP1 active site. SLiMs-mediated dephosphorylation are most impactful at low substrate concentrations and works most efficiently at low micromolar affinities as a compromise between the efficiency of targeting and the need to avoid product inhibition.

## Discussion

Our understanding of the specificity of dephosphorylation lags behind the knowledge of phosphorylation events, which is in part because the PPP family specificity relies on distal sequence motifs from the phospho-site and non-covalent protein interactions. It is also more difficult to investigate the phosphatase than kinases *in vitro* since it requires high-quality phospho-protein containing IDRs – samples that are usually challenging to prepare. Therefore, most biochemical studies of phosphatases are conducted using short phospho-peptides that are inherently unable to mimic the supra-molecular context of the complexes in which phosphatases work naturally.

The catalytic subunits of PPPs form holo-enzymes through non-covalent interactions with regulatory proteins that control the activity, sub-cellular localization, and local abundance of substrates. Recent proteomic experiments showed overlap between interactors and substrates of PP1 [3] and that many proteins regarded as regulators of the phosphatase act also as substrates for it [2]. Dephosphorylation thus often takes place in complexes where the active site and the phospho-site are physically linked via protein-protein interactions, where RVxF motif might be located thousands of residues from the phospho-site [2,3]. Each complex is unique, but the physical connection enhances the rate of encounters to an extent dependent on the supramolecular structure of the complex. In folded proteins, residues that are far apart in the sequence may be close in space. Therefore, length dependence of the spacing observed here only applies to intrinsically disordered regions and are not directly comparable to distances between SLiMs and phospho-sites observed in proteomic experiments. Similarly, as the docking interactions are distal to the phospho-site it will likely not change affinity upon dephosphorylation, which makes product inhibition likely for high affinity interactions. Neither the effect of the affinity nor the spacing is currently well described for phosphatases.

We sought to develop a model substrate that could model the supra-molecular context of phosphatase complexes yet convenient enough to allow kinetic experiments of many variants. We converged at a substrate with several key features: A phospho-site recognized both by PP1 and a convenient kinase (PKA), a protein background without off-target phosphorylation, two terminal purification tags to remove truncated substrates and a solubility tag that allows purification of variants with long IDRs. In combination, these features allow collection of high-quality kinetics for series of variants for theoretical modelling and present a system that can be adapted for other research questions.

The model system contains a few limitations: The GS-repeats used as spacers are a first order approximation of natural IDRs. Non-repetitive IDRs have sequence-specific effects on compaction that modulate effective concentrations in combination with length [25]. By removing sequence-specific effects the simplified IDRs allowed a clearer determination of linker length dependence than what would have been possible in any natural IDR. Additionally, residues flanking SLiMs interaction with the binding partner and modulate affinity [26,27], whereas our model assumes that the linker is inert and non-interacting. Titration of substrates without SLiMs shows partial competition at high concentrations (Fig. 4C) indicative of non-specific interactions. No protein is truly inert, and the malleability of IDRs make them prone to non-specific interactions. Such non-specific effects are hard to exclude when working with low affinity interactions, and provide a complication not included in the theoretical models. This difference might contribute to the discrepancy observed between experiments and models for weak SLiMs. Finally, we needed to combine two different substrate series to get a sufficient affinity range to sample both sides of the apex. There are slight differences between the binding interfaces of the NIPP1 and RVAF-series, for example the slightly extended binding site and bound-state heterogeneity of NIPP1 and slightly reduced C_eff_ value. However, flexible linkers ensure that catalytic efficiency is relatively insensitive to small changes in the complex structure.

While we only investigate PP1, we think the overall model for SLiM mediated dephosphorylation is likely to generalize to other PPPs that often recognize substrates via SLiMs. The model only relies on general physical principles such as an effective concentration of tethered substrate, and binding kinetics of the docking interaction. In many cases, the physical connection between active site and phospho-site will involve several protein chains and non-covalent interactions. In such cases, the weakest interaction will be rate limiting for both substrate targeting and product inhibition. Effective concentrations can be modelled for any system where a conformational ensemble can be generated – including combinations of folded disordered segments [28]. Proteomics experiments suggest that SLiMs can enhance dephosphorylation thousands of residues away, in agreement with theoretical modeling that suggest that effective concentrations decay relatively slowly with chain length. This suggests that the upper limit for SLiM to phosho-site linker in Fig. 2E may be due to non-specific interactions of the substrates with long GS repeats.

Recently, synthetic protein regulators of phosphatases have been developed including phosphatase-targeting chimeras that enhance dephosphorylation of specific targets [29,30]. Our data show that a small change in affinity can convert a regulatory protein from an enhancer to an inhibitor. The model presented here presents design principles that can guide future development of PhosTACs as well as for deciphering the effects of natural phosphatase regulators.

## Acknowledgement

This work was supported by grants to Magnus Kjærgaard from Carlsberg Foundation (CF22-0734), the Danish National Research Foundation (DNRF133), and instrument grants from the Carlsberg Foundation (CF24-1770). The “Biophysics and Biochemistry core facility” at Department of Molecular Biology & Genetics are thanked for technical support and instrument access.

The authors thank Maria Correia Davis for providing purified PKA, Daniel Erik Otzen for access to the stopped-flow instrument, Mette Hoffmann Asmussen and Jan Stanislaw Nowak for technical support, and Vili Petteri Lampinen, Alain A.M. André and Maria Correia Davis for critical reading and reviewing the manuscript.

## Materials and Methods

### Preparation of DNA constructs

PP1α_7-300_ plasmid was cloned into an in-house vector (RP1B), obtained from Addgene (no. 26566) [15]. The pGro7 plasmid was sourced from Takara #3340. Plasmids encoding substrates with C-terminal strep-tag were de novo synthesized by Twist Bioscience and codon-optimized for expression in *Escherichia coli*. The coding regions were cloned into pET28a(+) vectors using restriction sites Ndel and Xhol.

### Protein expression and purification

PP1α_7-300_ plasmid was co-transformed with pGro7 plasmid, which encodes the GroEL-ES chaperone into *BL21(DE3) E*.*coli* cell. The expression was performed following the protocol described by Kelker et al [15].

Plasmids encoding the substrate constructs were transformed into *BL21(DE3) E*.*coli* cells. Expression was carried out in TB medium supplemented with 50 µg/mL kanamycin. The culture was incubated at 37ºC and induced with 0.5 mM IPTG at an OD_600_ of ∼0.7, followed by incubation at 30ºC for 20 hours.

Cells were harvested by centrifugation (11,000 × g, 4ºC, 15 min). The cell pellets containing overexpressed PP1 were resuspended in lysis buffer (50 mM Tris-HCl, 700 mM NaCl pH 8.0), containing 5 mM imidazole, 1 mM MnCl_2_, 0.1% (v/v) Triton X-100, 0.2 mM PMSF, 1 µg/mL each of pepstatin, leupeptin and chymostatin. Cell pellets containing substrates were lysed in lysis buffer (50 mM Tris-HCl, 150 mM NaCl pH 7.6), supplemented with 0.2 mM PMSF, 1 µg/mL each of pepstatin, leupeptin and chymostatin. Cells were lysed using an ultrasonic homogenizer at 35% amplitude for 10 minutes (5 s on, 10 s off) while kept on ice. The lysates were centrifuged (22,000 × g, 4°C, 20 min), and the supernatants were applied to gravity flow columns packed with Ni-NTA resin (Qiagen). The columns were equilibrated and washed with their respective lysis buffers containing increasing concentrations of imidazole (5, 10, 20, 250 mM).

Further purification of PP1 was performed by size-exclusion chromatography (SEC) in TBS buffer (50 mM Tris-HCl, 700 mM NaCl, pH 8.0) using a Superdex 75 Increase column (GE Healthcare) at a flow rate 0.4 mL/min. Purified PP1 was stored in TBS pH 7.6 with 1 mM MnCl_2_ and 20% glycerol at –80ºC. Meanwhile, Ni-NTA chromatography eluate of the substrates were further purified by Strep-Tactin XT column. Bound substrates were eluted with Strep-elution buffer (50 mM Tris-HCl, 150 mM NaCl, 50 mM biotin, pH 7.6). Purified substrates were pooled and subsequently phosphorylated.

### Phosphorylation of the substrates

Protein kinase A (PKA) was used to phosphorylate the substrates, and the phosphorylation was assessed using SuperSep Phos-tag gel (50 µM, Fujifilm Wako). Phosphorylation was performed in TBS (50 mM Tris-HCl, 150 mM NaCl pH 7.6) supplemented with 10 mM magnesium acetate. Purified substrates at known concentration were pre-heated at 30ºC with 350 rpm shaking for 30 minutes. The reaction was initiated by adding 1 mM ATP (in excess relative to the substrate concentration) and 50 nM PKA and allowed to proceed for 2 hours. Phosphorylation was confirmed by the presence of bands corresponding to phosphorylated species, which migrate slower than the unphosphorylated control. Loading buffer for Phos-tag gel, composed of bromophenol blue (BPB), sodium dodecyl sulfate (SDS), glycerol, 0.5 M Tris-HCl pH 6.8, and 2-mercaptoethanol were added to the samples before loading onto the gel. The samples were run with Phos-tag running buffer (3 g Tris base, 1 g SDS, 72 g glycine in 1 L Mili-Q water) at 30 mA, 60V constant current for ∼1.5 h. Following phosphorylation, PKA was deactivated by heating up the solution to 90ºC for 5 minutes and residual ATP and ADP were removed by dialysis or using a CentriPure 25 (EMP Biotech) desalting column.

### PP1 activity assay

The activity freshly purified PP1 was determined using a 96-well plate (clear flat bottom, Greiner) with pNPP substrate. Each well contained 20 µL of 1 µM PP1 and 80 µL of pNPP substrate, with final concentrations of 0.5 mM, 1 mM, 2 mM, 4 mM, and 8 mM in 150 mM Bis-Tris buffer pH 6.5, 150 mM NaCl. The pNPP solution was incubated at 30°C for 1 minute with shaking before adding PP1. Absorbance was continuously measured at 405 nm, 30°C, every minute for 5 minutes. Kinetic measurements were carried out in quadruplicate for each substrate concentration, and the activity was calculated using:

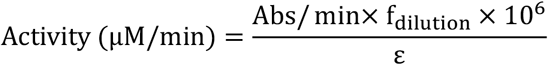

where Abs/min is appointed from the slope of kinetics in every substrate concentration, f_dilution_ is dilution factor, in this case is 5 (20 µL enzyme solution in 100 µL total reaction solution), 10^6^ as conversion from M to µM, ε is molar extinction coefficient of pNPP at 405 nm = 18000 M^-1^cm^-1^.

### Substrate dephosphorylation assay

Phosphate release from phosphorylated substrates were measured in SpectraMax Microplate Readers (Molecular Devices) on 96-well plate (clear flat bottom, Greiner) based on colorimetry from phosphate-ammonium-malachite green complex formation (green, max absorption at 620 nm). Malachite reagent mixture was prepared by mixing 1.32 mM malachite green (Thermo Scientific) in 10 mL concentrated H_2_SO_4_ and 50 mL deionized water; 30 mM H_2_SO_4_; ammonium molybdate (7.5% w/v, Thermo Scientific) in deionized water; and 11% Tween-20 with ratio 50:40:5:1 respectively and stepwise on ice. Substrates were adjusted to final concentrations of 10, 20, 30, 40, 60 and 80 µM in TBS (50 mM Tris-HCl, 150 mM NaCl, pH 7.6) supplemented with 1 mM MnCl_2_. The substrate solutions were pre-heated for 10 minutes at 30°C with shaking at 350 rpm. Dephosphorylation was initiated by adding 10 nM of PP1 to substrate solution in the reaction tube. 80 µL samples were taken every minute for 5 minutes and mixed with 20 µL of the malachite green reagent mixture to stop the reaction and convert the free phosphate. The dephosphorylation reaction was performed in triplicate. To ensure linearity, the product formation is limited to 20% of substrate consumption with a minimum of 3 data points.

Phosphate release rate of each substrate variant was plotted using specificity constant k_SP_, which equivalent to catalytic efficiency (k_cat_/K_M_). It was constructed by fitting the phosphate release as a function of substrate concentration, with following equation [16].

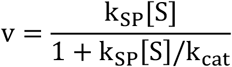

where v is the enzymatic rate, k_SP_ is the specificity constant, k_cat_ is the catalytic rate (V_max_/[E]), and [S] is the substrate concentration. All fittings and parameters were performed in GraphPad Prism 10 software.

### Fluorescence polarization (FP) direct binding assay

15-residue FITC-labelled NIPP1 (AKNKRVTFSEDDEII) prepared by GenScript (HPLC purified, 95% purity) was used as ligand for direct FP measurement. All measurements were performed on CLARIOstar Plus Multi-mode plate reader (BMG LabTech) using 384-well microplate reader (black, small-volume, non-binding Greiner) with 20 µL assay solution per well. The assay was conducted using 480 nm excitation and 535 nm emission filters. In the FP direct binding experiments, 10 nM FITC-labeled NIPP1 peptide was titrated by increasing concentration of PP1 in TBS (50 mM Tris-HCl, 150 mM NaCl, pH 7.6) and 0.05% Tween20. FP values were plotted as a function of log of PP1 concentrations. The dissociation constant K_d_ was obtained from following equation.

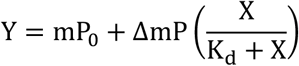

Where Y is mP value in each point, mP_0_ is initial mP value of the ligand, ΔmP is the difference between initial mP and end-pint mP, X is PP1 concentration, and K_d_ is dissociation constant. All fittings and parameters were performed in GraphPad Prism 10 software.

### Fluorescence polarization competition assay

FP competitive binding assay were measured using 30 nM FITC-labeled NIPP1 peptide and 250 nM of PP1, titrated with increasing concentration of unlabeled NIPP1(T202A) peptide (GenScript, HPLC purified, 95% purity) and full-length unphosphorylated substrates of other motives. The mixture was incubated for 30 minutes at room temperature prior to FP measurement. The IC_50_ values were calculated in GraphPad Prism 10 software following this equation.

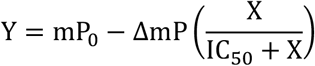

where Y is measured mP value, X is substrate/peptide concentration, mP_0_ is the highest mP value when substrate/peptide concentration is 0, ΔmP is the difference between mP_0_ and mP_final_. IC_50_ is the substrate/peptide concentration at the middle mP value. No constraint was applied to the substrates which reached binding saturation. However, mP_0_ and ΔP values for substrates did not show a binding saturation were constrained to 72 and 33 respectively.

The K_i_ or inhibition constant that translates to the K_d_ of substrate/peptide tested, is then calculated using this equation [31].

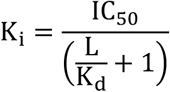

where L is ligand concentration (FITC-labeled NIPP1 peptide), K_d_ is dissociation constant of FITC-labeled NIPP1 peptide from FP direct binding assay, K_i_ is inhibition constant of the unlabeled-peptide which equivalent to the K_d_ of respected peptide motifs to PP1.

### Stopped-flow kinetics

Stopped-flow measurements were used to determine binding kinetics of NIPP1 to PP1. Dansyl-labelled 15-residue of NIPP1 peptide was obtained from GenScript (HPLC purified, 95% purity) as ligand. All measurements were performed on a Chirasan spectrometer equipped with Hg-Xe lamp and an Applied Photophysics stopped-flow mixer. Experiments were carried out in TBS (50 mM Tris-HCl, 150 mM NaCl, pH 7.6) at 30ºC. Fluorescence was monitored at 334 nm excitation (10 nm bandwidth) and 495 nm emission. To determine the association rate constant (k_on_), 100 nM dansyl-NIPP1 was mixed 1:1 with PP1 at various concentrations from 2.5 to 15 µM. Measurements were performed in triplicate over 5 s with 5000 time points and pressure hold enabled. Fluorescence traces were fitted to a single-exponential model between 0.1 and 1 s to obtain observed rate constant (k_obs_). The k_obs_ values were plotted against PP1 concentration and k_on_ was determined from the slope of the linear fit.

To determine the dissociation rate constant (k_off_), 100 nM dansyl-NIPP1 was preincubated with 4 µM PP1 and mixed 1:1 with 10 µM of NIPP1^196-210^T202A peptide as a competing ligand. Measurement was performed in triplicate over 100 s with 200 time points and pressure hold disabled. The fluorescence traces were fitted over the initial phase (0 – 3 s), and the resulting rate constant was taken as k_off_.

### Numerical simulations

Phosphate release rates as a function of SLiM dissociation rate (k_off_) was simulated in KinTek Global Kinetic Explorer [32] using the schematic pathways shown in Fig. 5A and Fig. S6A. The association rate constant (k_on_) was obtained from stopped-flow kinetic measurement and fixed at 2.4 × 10^6^ M^-1^s^-1^, while k_off_ was varied from 10^−2^ to 10^6^ s^-1^. The substrate association rate constant (k_1_) was set to a typical SLiM interaction of 10^7^ M^-1^s^-1^ [24] and the dissociation rate constant (k_-1_) was derived from the Michaelis constant (K_M_), yielding to 7340 s^-1^. The catalytic rate constant (k_2_) was set to the k_cat_ of untethered kemptide reaction, at 17.2 s^-1^. The product release rate constant (k_3_) was estimated at 1000 s^-1^ to favor for product release. The effective concentrations (C_eff_) of RVAF- and NIPP1-motif substrates were 6.54 mM and 6.12 mM respectively, corresponding to their spacers from core motif to kemptide. Simulations were performed at two substrate concentrations of 10 and 80 µM, which were the experimentally tested lowest and highest substrate concentrations. A constant enzyme concentration at 10 nM was also set to match the experimental part. The simulation time was adjusted to maintain less than 10% substrate conversion and ensure the linearity of the initial rate. The resulting slopes (b) were plotted as function of k_off_ using GraphPad Prism 10.

