## Supplemental information for "Affinity and distance dependence of SLiM-mediated dephosphorylation by protein phosphatase 1"

### Contents:

Protein and peptide sequences 3
Supplementary figures 5
Fig. S1. Purified proteins assessed by SDS-PAGE 5
Fig. S2. Substrate phosphorylation assessed by Phos-tag gel 6
Fig. S3. Malachite green assay 7
Fig. S4A. Measurement of in vitro phosphate release rates of modified RVAF_kemptide substrates 8
Fig. S4B. Measurement of in vitro phosphate release rates of modified single and composite SLiM_kemptide substrates 9
Fig. S5. Fluorescence polarization measurement of all substrates and PP1 10
Fig. S6. Measurement of in vitro phosphate release rates of modified NIPP1_kemptide substrates 11
Fig. S7. Association rate constant of NIPP1 measured by stopped-flow kinetics 12
Fig. S8. Productive possible ternary and quaternary complex formation 13
Supplementary tables 15
Table S1. Affinity of modified SLiMs. 15
Table S2. Rate constant for numerical simulation of steady-state phosphate release 16
Reference 17

#### Protein and peptide sequences

Color coding:

- 6xHis-tag
- Strep-tag
- Trx-1
- SLiMs: RVxF, SILK, MyPhoNE, ΦΦ
- PKA phosphorylation substrate motif, phospho-site is shown in bold

### PP1

MGSDKIHHHHHHENFLYFQGHMGSLNLDSIIGRLLEVQGSRPGKNVQLTENEIRGLCLKSREIFLSQP
ILLELEAPLKICGDIHGQYYDLLRLFEYGGFPPESNYLFLGDYVDRGKQSLETICLLLAYKIKYPENF
FLLRGNHECASINRIYGFYDECKRRYNIKLWKTFTDCFNCLPIAAIVDEKIFCCHGGLSPDLQSMEQI
RRIMRPTDVPDQGLLCDLLWSDPDKDVQGWGENDRGVSFTFGAEVVAKFLHKHDLDLICRAHQVVEDG
YEFFAKRQLVTLFSAPNYCGEFDNAGAMMSVDETLMCSFQILKPAD

##### PKA

MGSSHHHHHHSSGLVPRGSHMVTDEDIRKQEERAQQVRKKLEEALMADASQSGTGNAAAAKKGSEQES
VKEFLAKAKEDFLKKWETPSQNTAQLDQFDRIKTLGTGSFGRVMLVKHKESGNHYAMKILDKQKVVKL
KQIEHTLNEKRILQAVNFPFLVKLEFSFKDNSNLYMVMEYVAGGEMFSHLRRIGRFSEPHARFYAAQI
VLTFEYLHSLDLIYRDLKPENLLIDQQGYIQVTDFGFAKRVKGRTWTLCGTPEYLAPEIILSKGYNKA
VDWWALGVLIYEMAAGYPPFFADQPIQIYEKIVSGKVRFPSHFSSDLKDLLRNLLQVDLTKRFGNLKN
GVNDIKNHKWFATTDWIAIYQRKVEAPFIPKFKGPGDTSNFDDYEEEEIRVSINEKCGKEFTEF

##### RVxF_kemptide substrates

MGSSHHHHHHSSGLVPRGSHMMSDKIIHLTDDSFDTDVLKADGAILVDFWAEWCGPCKMIAPILDEIA
DEYQGKLTVAKLNIDQPGTAPKYGIRGIPTLLLFKNGEVAATKVGALSKGQLKEFLDANLALVPRGSR
VAF(GS)$_{n}$LRRA**S**LGWSHPQFEK

##### RVxF_LRRAALG substrate

MGSSHHHHHHSSGLVPRGSHMMSDKIIHLTDDSFDTDVLKADGAILVDFWAEWCGPCKMIAPILDEIA
DEYQGKLTVAKLNIDQPGTAPKYGIRGIPTLLLFKNGEVAATKVGALSKGQLKEFLDANLALVPRGSR
VAF(GS)$_{10}$LRRA**A**LGWSHPQFEK

##### RAAA_kemptide substrate

MGSSHHHHHHSSGLVPRGSHMMSDKIIHLTDDSFDTDVLKADGAILVDFWAEWCGPCKMIAPILDEIA
DEYQGKLTVAKLNIDQPGTAPKYGIRGIPTLLLFKNGEVAATKVGALSKGQLKEFLDANLALVPRGSR
AAA(GS)$_{10}$LRRA**S**LGWSHPQFEK

##### NonSLiM_kemptide control

MGSSHHHHHHSSGLVPRGSHMMSDKIIHLTDDSFDTDVLKADGAILVDFWAEWCGPCKMIAPILDEIA
DEYQGKLTVAKLNIDQPGTAPKYGIRGIPTLLLFKNGEVAATKVGALSKGQLKEFLDANLALVPRGS(
GS)$_{12}$LRRA**S**LGWSHPQFEK

##### SILK_kemptide substrates

MGSSHHHHHHSSGLVPRGSHMMSDKIIHLTDDSFDTDVLKADGAILVDFWAEWCGPCKMIAPILDEIA
DEYQGKLTVAKLNIDQPGTAPKYGIRGIPTLLLFKNGEVAATKVGALSKGQLKEFLDANLALVPRGSG
SSILK(GS)$_{n}$LRRA**S**LGWSHPQFEK

#### MyPhoNE_kemptide substrates

MGSSHHHHHHSSGLVPRGSHMMSDKIIHLTDDSFDTDVLKADGAILVDFWAEWCGPCKMIAPILDEIA
DEYQGKLTVAKLNIDQPGTAPKYGIRGIPTLLLFKNGEVAATKVGALSKGQLKEFLDANLALVPRGSR
GDQVKGW$(GS)_n$LRRA**S**LGWSHPQFEK

#### ΦΦ_kemptide substrates

MGSSHHHHHHSSGLVPRGSHMMSDKIIHLTDDSFDTDVLKADGAILVDFWAEWCGPCKMIAPILDEIA
DEYQGKLTVAKLNIDQPGTAPKYGIRGIPTLLLFKNGEVAATKVGALSKGQLKEFLDANLALVPRGSA
F$(GS)_n$LRRA**S**LGWSHPQFEK

#### NIPP1$^{196-210}$_kemptide substrate

MGSSHHHHHHSSGLVPRGSHMMSDKIIHLTDDSFDTDVLKADGAILVDFWAEWCGPCKMIAPILDEIA
DEYQGKLTVAKLNIDQPGTAPKYGIRGIPTLLLFKNGEVAATKVGALSKGQLKEFLDANLALVPRGS$^{19}$
$^{6}$AKNKRVTFSEDDEII$^{210}$$(GS)_{10}$LRRA**S**LGWSHPQFEK

#### Table of substrate and enzyme constructs used in this work:

| Enzyme | RVxF substrate | Modified SLiMs substrate | Control substrates |
|---|---|---|---|
| PP1 | RVxF_$(GS)_3$_kemptide | SILK_$(GS)_{10}$_kemptide | NoSLiM_kemptide |
| PKA | RVxF_$(GS)_6$_kemptide | SILK_$(GS)_{20}$_kemptide | RVxF_$(GS)_{10}$_LRRAALG |
| | RVxF_$(GS)_8$_kemptide | MyPhoNE_$(GS)_{10}$_kemptide | SILK_$(GS)_{10}$_ LRRAALG |
| | RVxF_$(GS)_{10}$_kemptide | MyPhoNE_$(GS)_{20}$_kemptide | ΦΦ_$(GS)_{10}$_ LRRAALG |
| | RVxF_$(GS)_{15}$_kemptide | ΦΦ_$(GS)_{10}$_kemptide | MyPhoNE_$(GS)_{10}$_ LRRAALG |
| | RVxF_$(GS)_{20}$_kemptide | ΦΦ_$(GS)_{20}$_kemptide | |
| | RVxF_$(GS)_{30}$_kemptide | SILK+RVAF_$(GS)_{10}$_kemptide* | |
| | RVxF_$(GS)_{40}$_kemptide | ΦΦ+RVAF_$(GS)_{10}$_kemptide* | |
| | RVxF_$(GS)_{50}$_kemptide | MyPhoNE+RVAF_$(GS)_{10}$_kemptide* | |
| | RAAA_$(GS)_{10}$_kemptide | SILK+NIPP1(T202A)_$(GS)_{10}$_kemptide* | |
| | | NIPP1(T202A)_kemptide | |
| | | NIPP1(T202Q)_kemptide | |
| | | NIPP1(K199S/T202A)_kemptide | |
| | | NIPP1(K199L/T202A)_kemptide | |
| | | NIPP1(V201A/T202A/F203A)_kemptide | |

*Distance between 2 combined SLiMs:

- SILK_NIPP1: $(GS)_5$
- SILK_RVAF: $(GS)_5$
- ΦΦ_RVAF: $(GS)_8$
- MyPhoNE_RVAF: $(GS)_7$

#### Peptide

- FITC-AKNKRVTFSEDDEII
- NIPP1$^{196-210}$T202A: AKNKRVAFSEDDEII
- Dansyl- AKNKRVTFSEDDEII

#### Supplementary figures

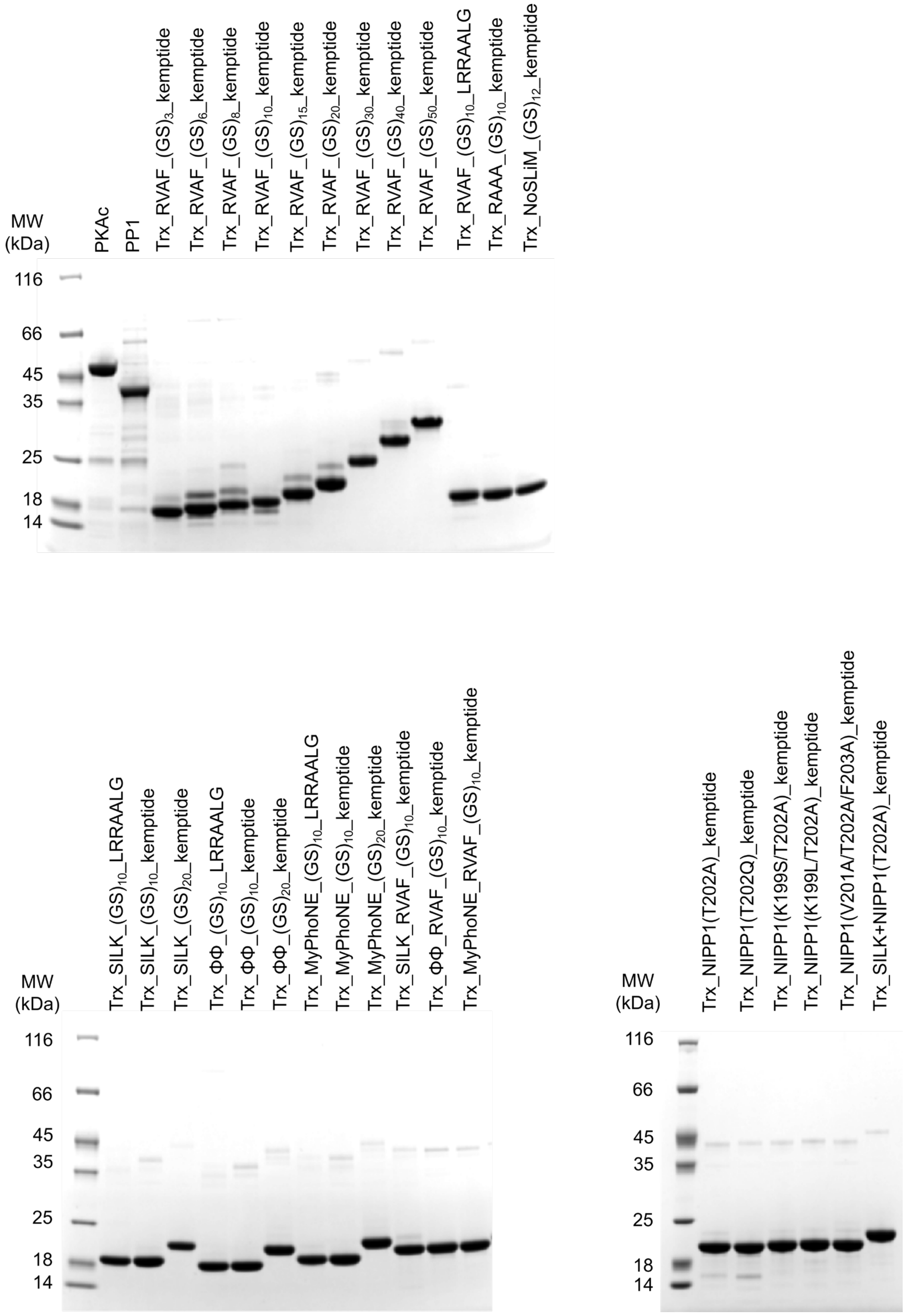


**Fig. S1. Purified proteins assessed by SDS-PAGE.** Gradient 4 – 20% polyacrylamide gel was used to assess all purified recombinant proteins: PKAc, PP1, RVAF substrates with various lengths, modified and composite SLiMs substrates, mutated NIPP1 substrates. 2µg of proteins are loaded to each lane and stained by Coomassie following the run (200 V, 45 minutes).

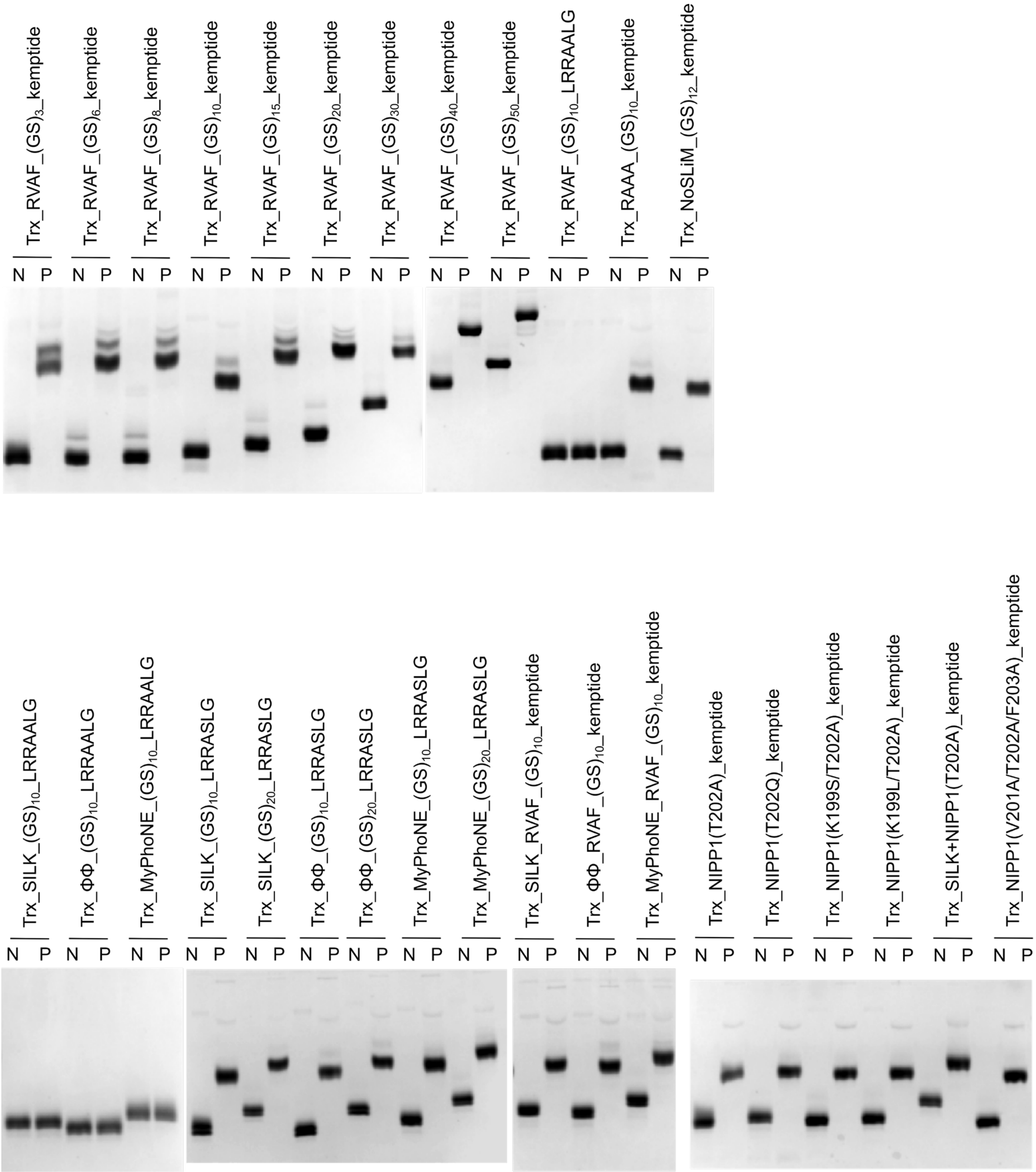


**Fig. S2. Substrate phosphorylation assessed by Phos-tag gel**. Phos-tag gel containing $Zn^{2+}$ ions reduces the migration speed of phosphorylated proteins, resulting in a higher band. 0.5 µg of non-phosphorylated protein (N) and phosphorylated (P) were loaded in each lane. All substrates got phosphorylated except mutated kemptide: left-hand gel with samples ending with LRRAALG sequence.

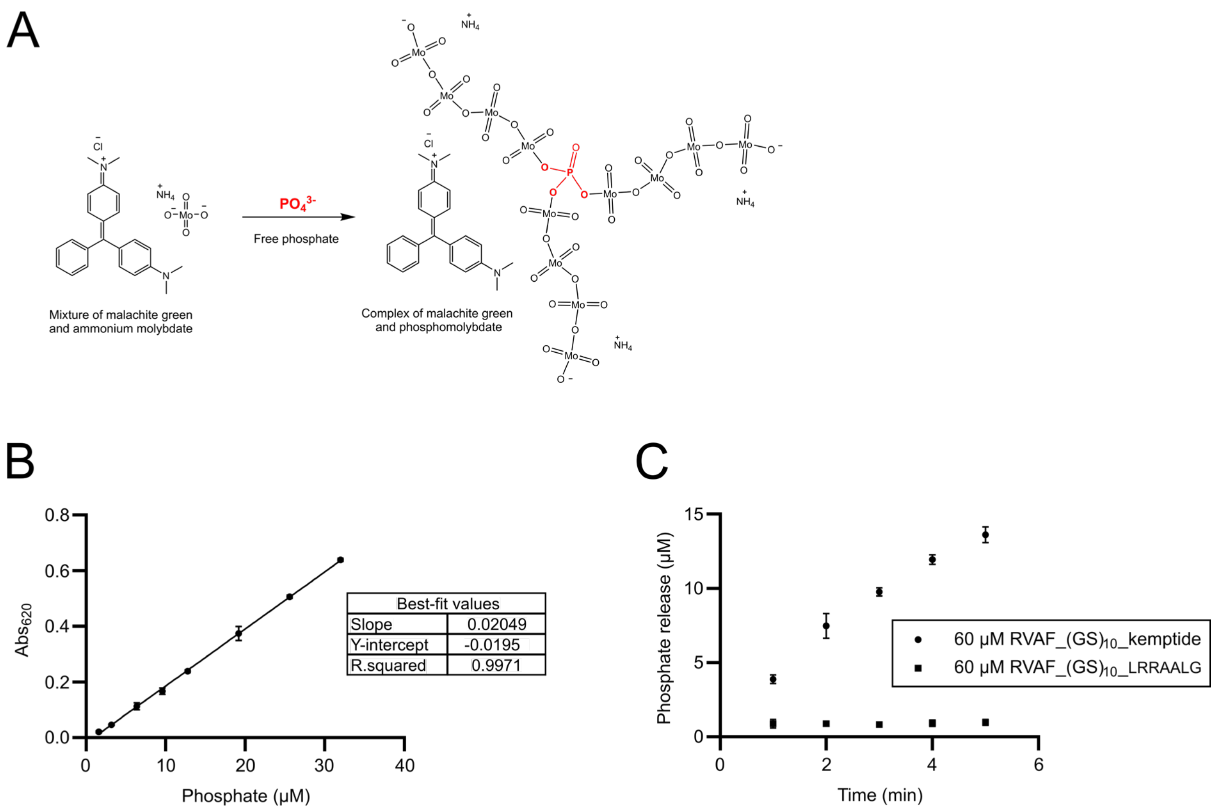


**Fig. S3. Malachite green assay. A)** Complex formation between malachite green, ammonium molybdate and inorganic phosphate forming stable green color, detectable with visible wavelength at 620 nm. **B)** Standard phosphate calibration curve using 1.6 – 32 μM phosphate standard solution. **C)** Phosphate release rate of kemptide (LRRASLG) substrate compared to mutated (LRRAALG) with same linker length as negative phosphorylation control at 60 μM substrate concentrations.

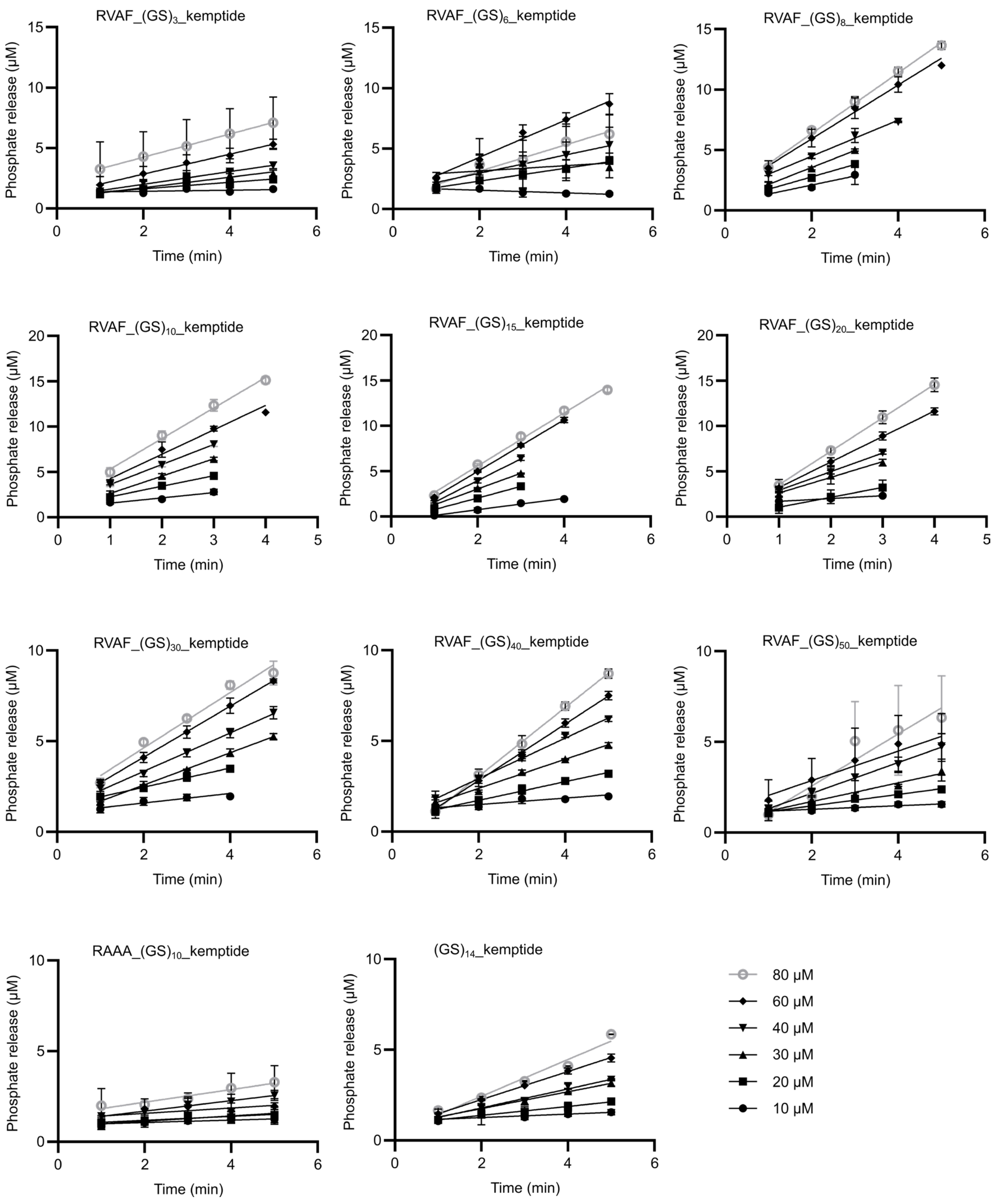


**Fig. S4A. Measurement of in vitro phosphate release rates of modified RVAF_kemptide substrates.** Phosphate release rate of RVAF_kemptide with various linker length, RAAA_kemptide, and NoSLiM_kemptide variants were tested at 10 – 80 μM substrate concentration and 10 nM PP1. The linear regressions were limited to 20% substrate consumption with minimum 3 points to fit.

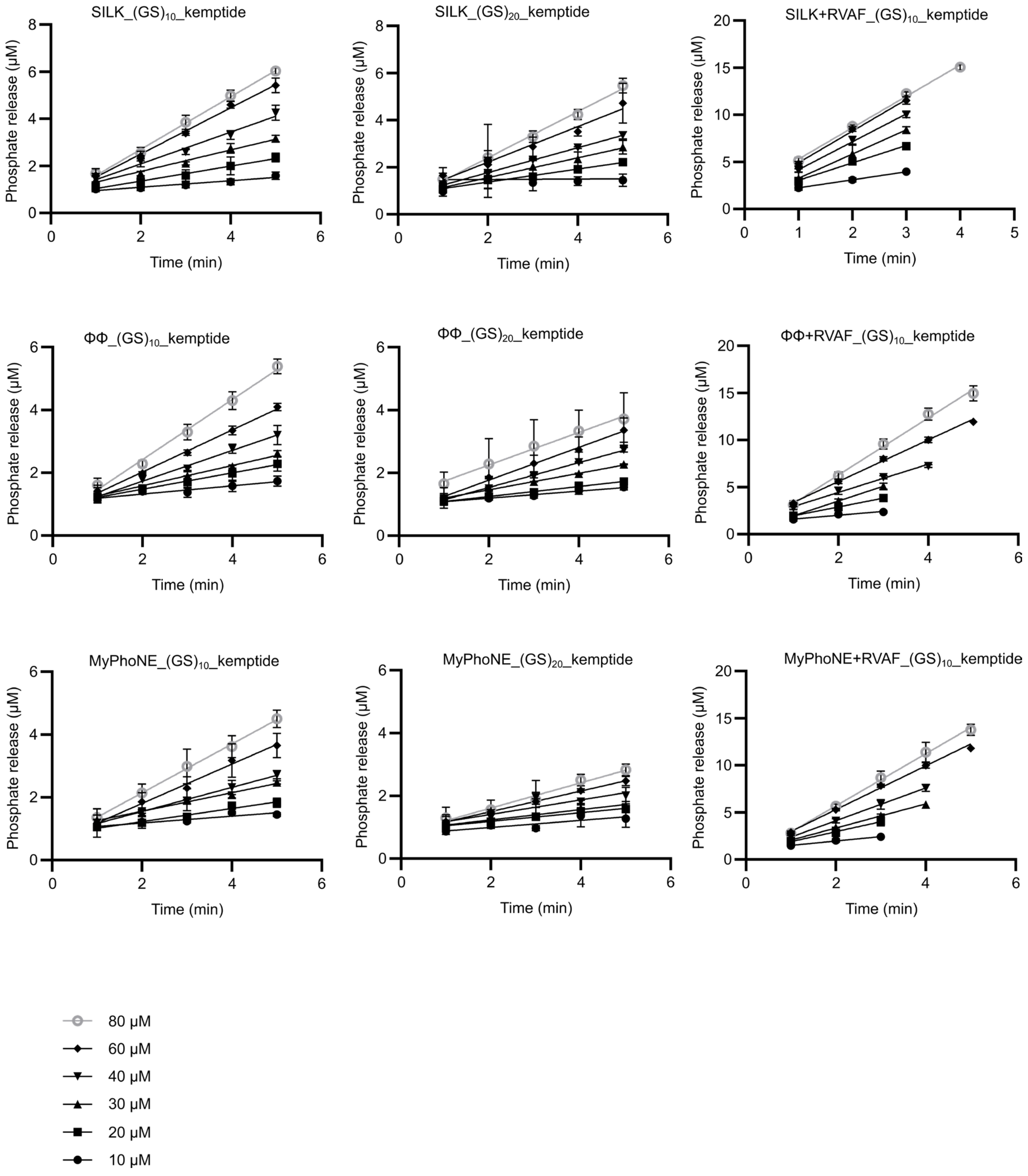


**Fig. S4B. Measurement of in vitro phosphate release rates of modified single and composite SLiM_kemptide substrates.** Phosphate release rate of single SLiM: SILK_kemptide, ΦΦ_kemptide and MyPhoNE_kemptide with 20- and 40-residue linkers and double SLiMs: SILK+RVAF_kemptide, ΦΦ+RVAF_kemptide and MyPhoNE+RVAF_kemptide variants were tested at 10 – 80 µM substrate concentration and 10 nM PP1. The linear regressions were limited to 20% substrate consumption with minimum 3 points to fit.

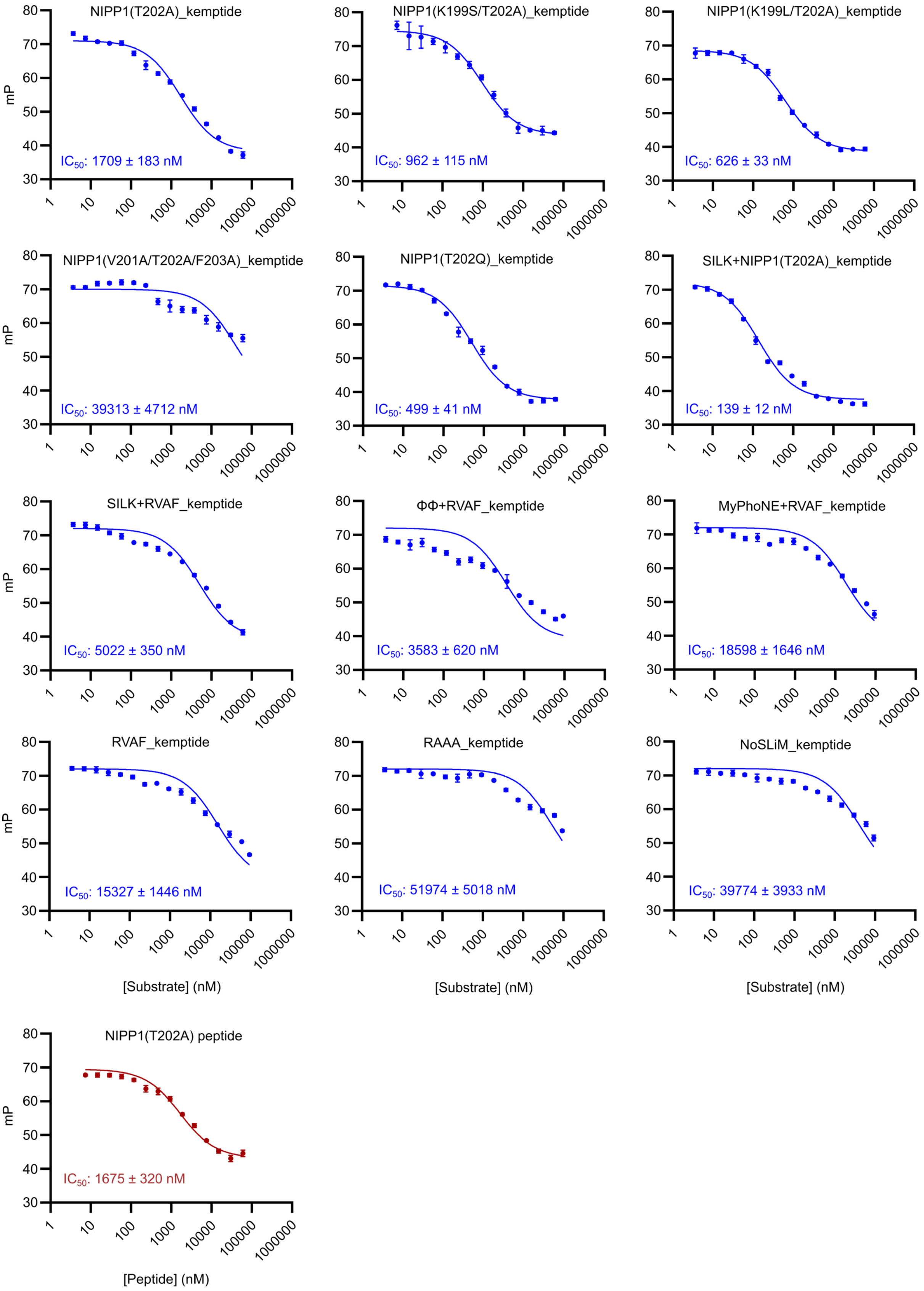


**Fig. S5. Fluorescence polarization measurement of all substrates and PP1.** Affinity measurement of full-length SLiM-modified substrates and the composites with PP1 using fluorescence polarization competition assay. Blue fits are full-length SLiM_kemptide, while red plot is 15-residue NIPP1(T202A) peptide. The midpoint of binding saturation shows $IC_{50}$ value.

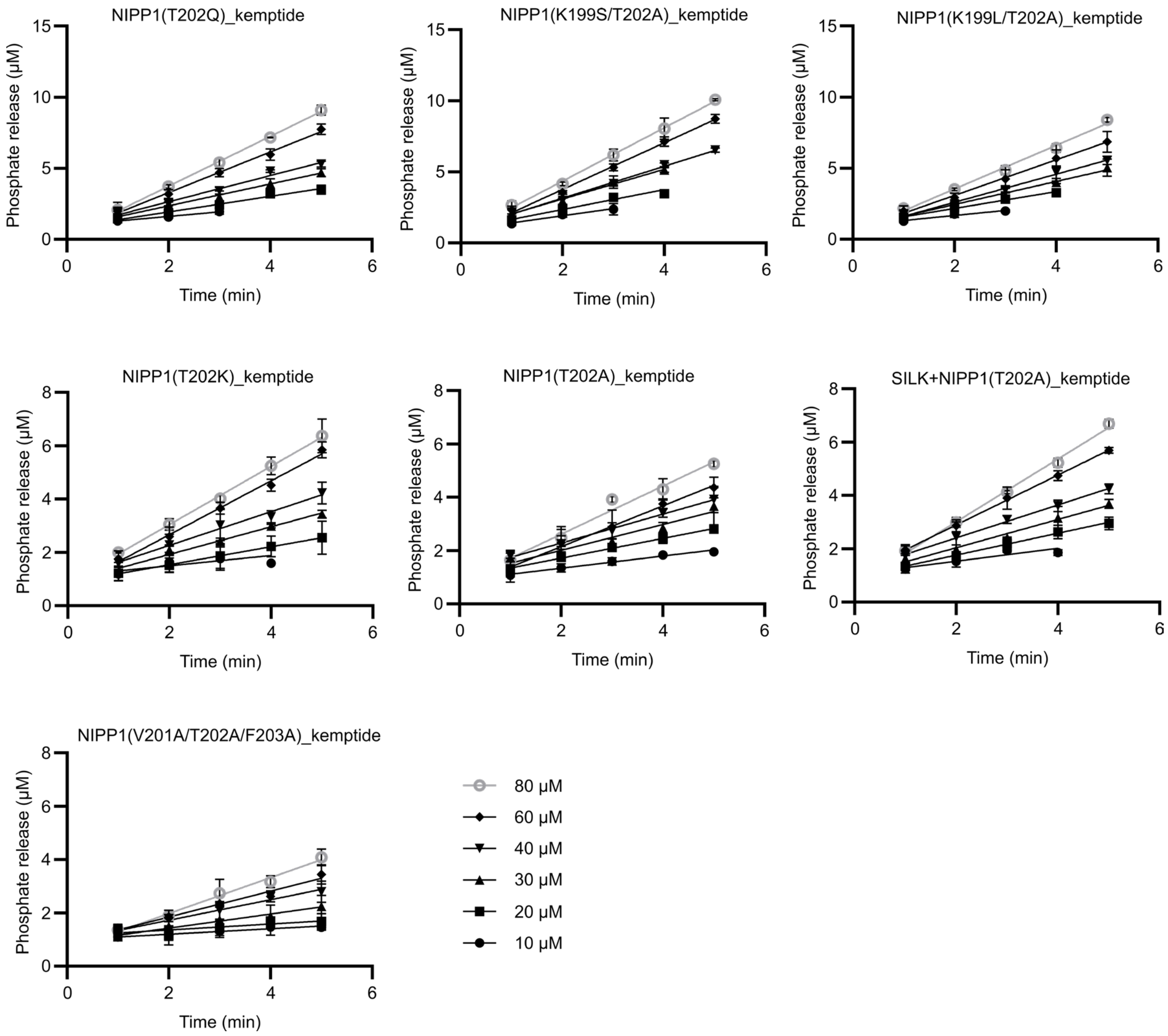


**Fig. S6. Measurement of in vitro phosphate release rates of modified NIPP1_kemptide substrates.** Phosphate release rate of six variants of NIPP1 ($^{196}$AKNKRVTFSEDDEII$^{210}$)_kemptide substrates and 1 composite form (+SILK) were tested at 10 – 80 µM substrate concentration and 10 nM PP1. The linear regressions were limited to 20% substrate consumption with a minimum of 3 points to fit.

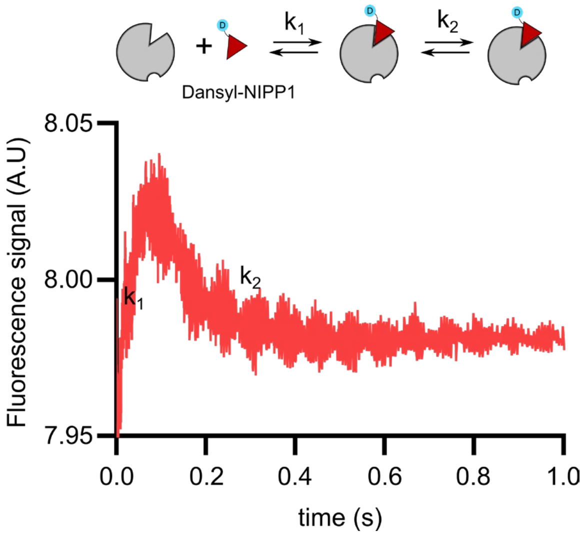


**Fig. S7. Association rate constant of NIPP1 measured by stopped-flow kinetics.** Schematic representation of PP1 and dansyl-labelled NIPP1 (ligand) association rates with two bound conformations ($k_1$ and $k_2$). Fluorescence traces recorded at 334 nm excitation and 495 nm emission using 12.5 µM PP1 with 1:1 ratio with 100 nM ligand (final concentration of 50 nM ligand and 6.25 µM PP1 upon mixing). Typical spectral recorded has biphasic rate constants.

A

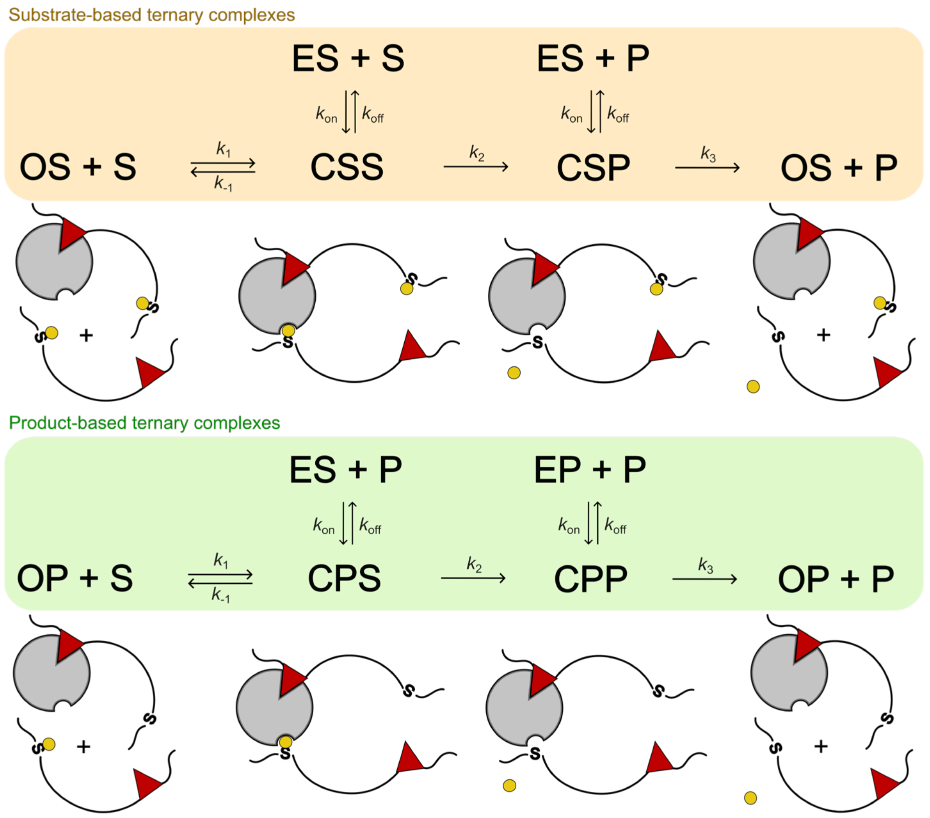


B

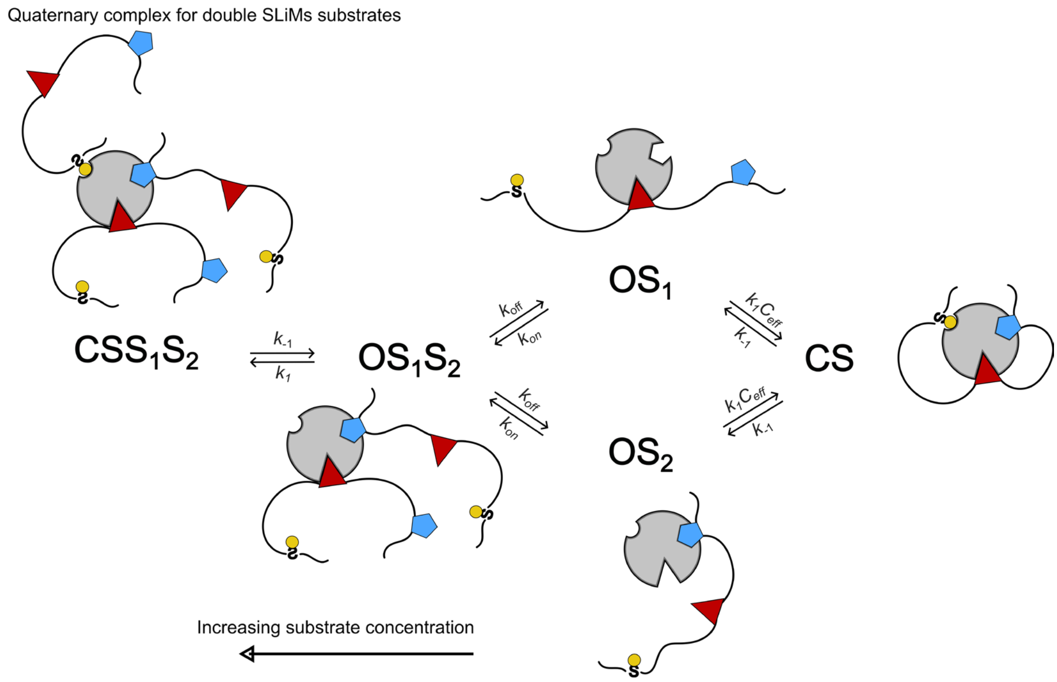


**Fig. S8. Productive possible ternary and quaternary complex formation. A)** Substrate- and product-based ternary complex between enzyme-substrate complex and the second substrate molecules arrive at the active site. This ternary pathway was adapted from Dyla,et al, 2022 [1]. Enzyme (E), substrate (S), and product (P) are shown. ES and EP indicate the enzyme's active site is bound with the substrate or product. OS and OP denote open state of enzyme-substrate or enzyme-product complex, while CSS/CSP and CPS/CPP represent closed conformation of the ternary complex. These two pathways are included in the numerical simulations. **B)** Possible quaternary complex formation for substrates with double SLiMs: SILK+RVAF, MyPhoNE+RVAF, ΦΦ+RVAF, SILK+NIPP1 variants at high concentrations. $S_1$ indicates substrate docks in RVxF binding pocket of PP1, while $S_2$ indicates substrate docks in the second docking pocket. These pathways/species are not included in the numerical simulations. All pathways favor product formation.

#### Supplementary tables

**Table S1. Affinity of modified SLiMs.**

| Motifs | $K_d$ (nM) | Measurement method |
|---|---|---|
| NIPP1 peptide | 252 ± 29 | Direct FP |
| NIPP1(T202A) peptide | 1523 ± 291 | Competitive FP |
| NIPP1(T202A)_kemptide | 1527 ± 164 | Competitive FP |
| NIPP1(K199S_T202A)_kemptide | 860 ± 103 | Competitive FP |
| NIPP1(K199L_T202A)_kemptide | 559 ± 29 | Competitive FP |
| NIPP1(V201A_T202A_F203A)_kemptide | 23280 ± 3129 | Competitive FP |
| NIPP1(T202Q)_kemptide | 446 ± 37 | Competitive FP |
| SILK+NIPP1(T202A)_kemptide | 125 ± 11 | Competitive FP |
| RVAF_kemptide | 13933 ± 1314 | Competitive FP |
| SILK+RVAF_kemptide | 4565 ± 318 | Competitive FP |
| ΦΦ+RVAF_kemptide | 3257 ± 564 | Competitive FP |
| MyPhoNE+RVAF_kemptide | 16907 ± 1496 | Competitive FP |
| RAAA_kemptide | 47249 ± 4562 | Competitive FP |
| NoSLiM_kemptide | 36158 ± 3575 | Competitive FP |

**Table S2. Rate constant for numerical simulation of steady-state phosphate release**

| Parameter | Value | Notes |
|---|---|---|
| [E] | 0.00000001 M | Experiment's enzyme concentration |
| [S] | 0.00001 M<br>0.00008 M | Experiment's lowest and highest substrate concentrations |
| $k_{on}$ | $3.3 \times 10^7$ $M^{-1}.s^{-1}$ | Calculated from stopped-flow experiment |
| $k_{off}$ | Varied | |
| $k_1$ | $1 \times 10^7$ $M^{-1}.s^{-1}$ | Typical average enzyme-SLiM association rate constant [1] |
| $k_{-1}$ | 7340 $s^{-1}$ | From $K_M$ = 0.000734 M of phosphate release kinetics |
| $k_2$ | 8.3 $s^{-1}$ | From average $k_{cat}$ of tethered reactions = 8.3 $s^{-1}$ |
| $k_3$ | 1000 | Estimation to not becoming rate-limiting step $k_3$>>$k_2$ |
| $C_{eff}$ (RVAF) | 0.0065406 M | From $C_{eff}$ calculator app [2] |
| $C_{eff}$ (NIPP1) | 0.0061167 M | |
| $K_{d\_apex}$ [S] = 10 µM<br>RVAF<br>NIPP1 | <br>6820 nM<br>6590 nM | From derivation of analytical equation [1] |
| $K_{d\_apex}$ [S] = 80 µM<br>RVAF<br>NIPP1 | <br>19300 nM<br>18600 nM | |

Reaction models and rate constant parameters included in the numerical simulation are available in Fig. 5 and Fig. S6A, adapted from Dyla, et al, 2022. Full derivative resulting in an equation to estimate $k_{off\_apex}$ is also adapted from the same article [1].

$$k_{off\_apex} = \sqrt{\frac{k_{cat}}{K_M} k_{on} C_{eff} S_0}$$

$$K_{d\_apex} = k_{off\_apex} \times k_{on}$$

$k_{off\_apex}$ : optimum tethering strength ($s^{-1}$)
$k_{cat}$ : catalytic constant
$K_M$ : Michaelis-Menten constant
$k_{on}$ : association rate constant
$C_{eff}$ : effective concentration
$S_0$ : substrate concentration
$K_{d\_apex}$ : optimum dissociation rate constant for tethering reaction (nM)
